# Collective Regulation of Metabolism, Development and Longevity Through a Socially Exchanged Fluid

**DOI:** 10.1101/2021.01.28.428590

**Authors:** SM Hakala, M-P Meurville, M Stumpe, AC LeBoeuf

**Affiliations:** Department of Biology, University of Fribourg, Chemin du Musée 10, 1700, Fribourg, Switzerland; Metabolomics and Proteomics Platform, Department of Biology, University of Fribourg, Chemin du Musée 10, 1700, Fribourg, Switzerland

**Author notes:** Corresponding author: Adria LeBoeuf.

**Keywords:** Aging, Hymenoptera: Formicidae, individual variation, life history evolution, machine learning, maturation, network, population proteomics, reproduction, senescence, social behavior, social evolution, social physiology, superorganism, tradeoff

## Abstract

Socially exchanged fluids, like seminal fluid and milk, allow individuals to molecularly influence conspecifics. Many social insects have a social circulatory system, where food and endogenously produced molecules are transferred mouth-to-mouth (stomodeal trophallaxis), connecting all the individuals in the society. To understand how these endogenous molecules relate to colony life, we used quantitative proteomics to investigate the trophallactic fluid within colonies of the carpenter ant *Camponotus floridanus*. We show that different stages of the colony life cycle circulate different types of proteins: young colonies prioritize direct carbohydrate processing; mature colonies prioritize accumulation and transmission of stored resources. Further, colonies circulate proteins implicated in oxidative stress, ageing, and social insect caste determination, potentially acting as superorganismal hormones. Brood-caring individuals that are also closer to the queen in the social network (nurses) showed higher abundance of oxidative stress-related proteins. Thus, trophallaxis behavior provides a mechanism for distributed metabolism in social insect societies.

## Introduction

In the course of social evolution, related organisms have formed cooperative entities such as multicellular organisms or groups of social animals (*1*–*3*). In social animal groups, collective decisions on movement, reproduction and even development are needed for survival (*4*, *5*). Countering this drive toward shared interests and collective good, conflicting interests complicate cooperation, but fundamentally also drive the evolution of increasingly coordinated behavior by driving the evolution of control mechanisms to suppress such social conflicts (*6*)(*6*–*8*). Some social groups have taken this coordination to a very high level: social insect societies develop and function as a single unit instead of as competing individuals, as ‘superorganisms’ paralleling the development of multicellular organisms as a single unit rather than as a set of competing cells (*9*, *10*).

In these superorganismal societies, reproductive queens and males function as the germline, and workers as the soma. Similarly to different tissues in multicellular organisms, workers can be further specialized and exhibit division of labor across different behavioral and morphological castes (*6*). While morphological castes are determined during development, the behavioral caste of an individual worker typically changes during its lifetime. At the beginning of their adult life, workers specialize inside the nest as nurses focusing on brood care, and as they age, they switch to foraging outside of the nest (*11*). Social insect colonies also go through life stages. Young colonies have an initial growth phase where they solely produce one type of worker, and only later in their life cycle they may produce more specialized worker castes and finally, males and queens (*12*). The switch to reproductive phase is a major life-history transition at the colony level, and connected to female caste determination. In social Hymenoptera, determination of whether a female larva develops into a queen or a worker, and what kind of worker exactly, is controlled by intricate differences of gene expression of the same female genome, guided primarily by environmental factors, in particular nutrition and social cues, but sometimes also influenced by genetics (*13*–*16*).

Coordinated function of tightly integrated groups such as social insect colonies, and subgroups such as their different castes, has been described as social physiology (*17*), consisting of various behavioral, morphological and molecular mechanisms that ensure cooperation and inclusive fitness benefits for all group members. As a part of their social physiology, some social insect societies have developed a form of social circulatory system, where nutrition and endogenously produced functional molecules, such as hormones, are transferred mouth-to-mouth from the foregut of one individual to another (*18*, *19*). This social fluid transfer is called stomodeal trophallaxis (*20*). It ensures not only that food is distributed to all adults and larvae within the colony, but also that all individuals of the colony are interconnected through shared bodily fluids. Trophallactic fluid of ants and bees typically contains endogenous proteins involved in digestion, immune defense and developmental regulation (*18*), indicating that this fluid transmits more than food.

Molecular signals are important in controlling the colony life histories and guiding caste determination both at the colony level and at the individual level. Queen pheromones are central signaling molecules acting across individuals (*21*–*23*). Juvenile hormone and vitellogenin are central signaling molecules in classical insect development that may also play across-individual roles in some social insects (*18*, *19*, *24*, *25*). Together with fundamental nutrient-response signaling pathways (insulin, TOR), these molecules establish the developmental trajectories of individuals (*26*, *27*). In solitary organisms, such molecules are produced and function solely within the organism’s own body. In contrast, in social Hymenoptera even the molecules traditionally functioning within-individuals can be secreted to the crop and distributed among the society members through trophallaxis and the social circulatory system (*18*).

Molecular components transmitted through trophallaxis, namely juvenile hormone and juvenile hormone esterase-like proteins, have been shown to influence the development of ant larvae (*18*, *19*). Thus it is possible that molecules in trophallactic fluid may influence caste determination, similarly to honeybee workers feeding larvae with royal jelly to direct their development toward a queen fate (*18*, *28*–*31*). The molecular functions of trophallactic fluid are still largely unstudied, but it is known, for example, that social isolation changes its composition (*18*), suggesting that this fluid correlates with social environment. Further, we propose that trophallactic fluid could both reflect and affect the social environments of the colony, thus providing important environmental cues for collective decision making. However, it is still largely unknown how much and what kind of qualitative and quantitative variation is present in trophallactic fluid and what causes it.

If indeed trophallactic fluid acts as a form of social circulatory system, managing distributed metabolic processes related to colony maturation, endogenously produced factors should correlate with colony life stages. To test this, we analyzed the trophallactic fluid proteome of the carpenter ant *Camponotus floridanus* at different scales, monitoring colonies under different biotic and abiotic conditions and in different individuals within colonies.

## Results

We sought to determine whether the endogenously produced proteins present in trophallactic fluid create a robust biomarker-like signature of colony status. To assess this, we analyzed the trophallactic fluid proteomes of colonies at different stages in the colony life-cycle (*Young vs. Mature*), of colonies in natural conditions or kept in the lab (*Field vs. Lab*), and between colonies found on different nearby islands (*East vs. West*) (Figure 1, SI Table 1). Because trophallactic fluid proteins may be differentially expressed, transmitted, and/or sequestered across the social network of a colony, we also analyzed trophallactic fluid proteomes of single individuals in different colony ‘tissues’ – in-nest workers taking care of brood and out-of-nest workers (*Nurse vs. Forager*).

**Figure 1:**
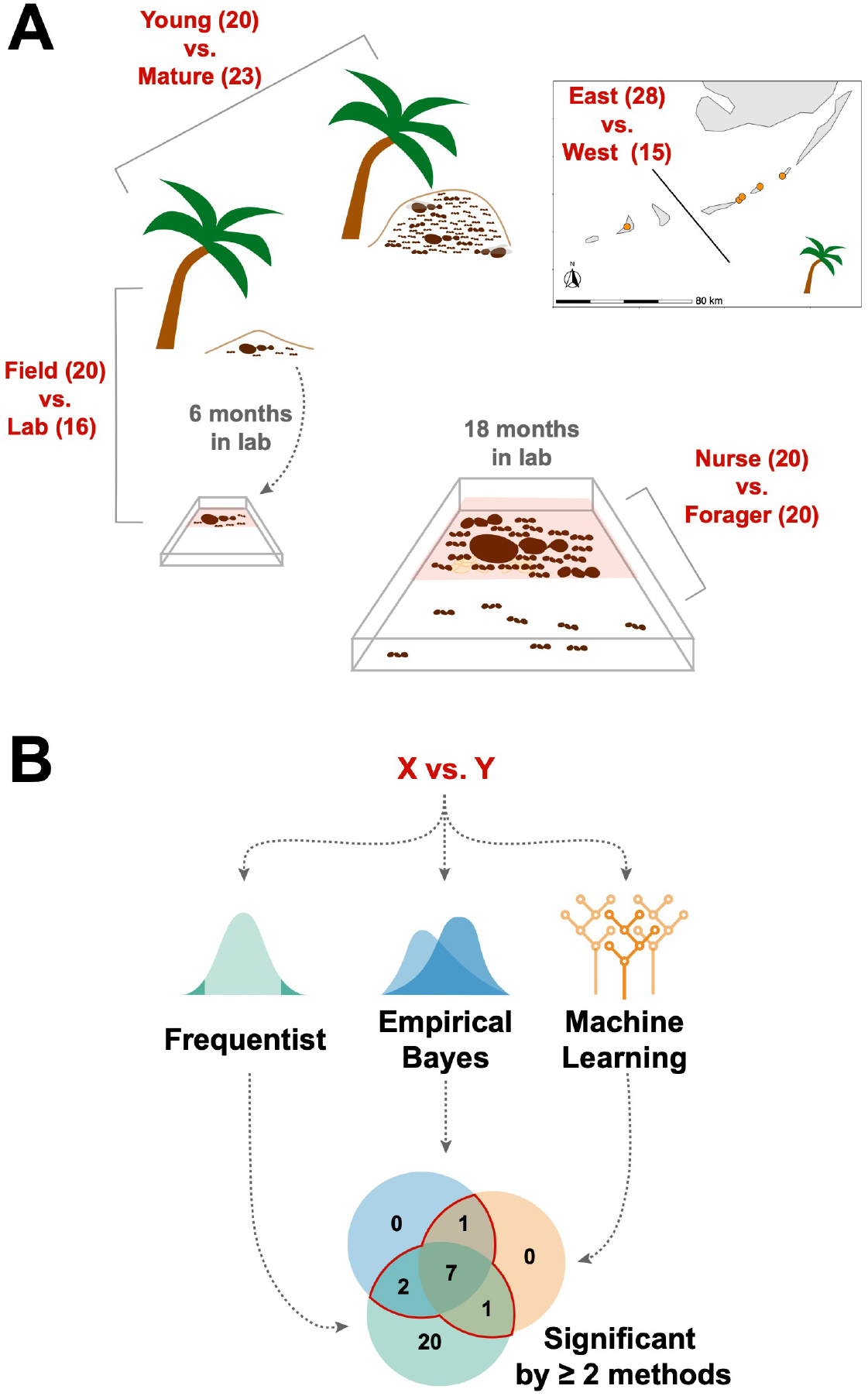
Schematic of study design. A. Four comparisons, *Young vs. Mature*, *Nurse vs. Forager*, *Field vs. Lab*, and *East vs. West*, analyzed in this study with sample numbers indicated in parentheses. In all comparisons sample numbers indicate colonies with the exception of *Nurse vs. Forager*, where samples are from single individuals, ten each from four colonies. Palm trees indicate field samples and boxes indicate laboratory samples. B. Schematic of analysis approach to find robustly differing proteins in each comparison.

### Overall proteome variation

Over the 73 colony samples and 40 single individual trophallactic fluid samples analyzed, a total of 519 proteins were identified (Figure 2). Trophallactic fluid samples contained a set of 27 ‘core’ trophallactic fluid proteins that were present in all samples regardless of life-cycle, life-stage or environmental conditions. Fifty-seven percent of the 519 proteins we observed were present in less than half of the samples. Even though the most common proteins displayed higher average abundance, across the entire dataset, protein abundance did not correlate with the proportion of samples containing the protein – even proteins present in only a small proportion of the samples in some cases exhibited high abundance (SI Figure 1). The overall protein abundance was higher in colony samples relative to single individual samples, reflective of the larger trophallactic fluid volume collected. The number of proteins identified for a given sample correlated with trophallactic fluid sample volume (Pearson correlation test p < 0.05, r = 0.24 for colony samples and p < 0.05, r = -0.40 for single individual samples).

**Figure 2:**
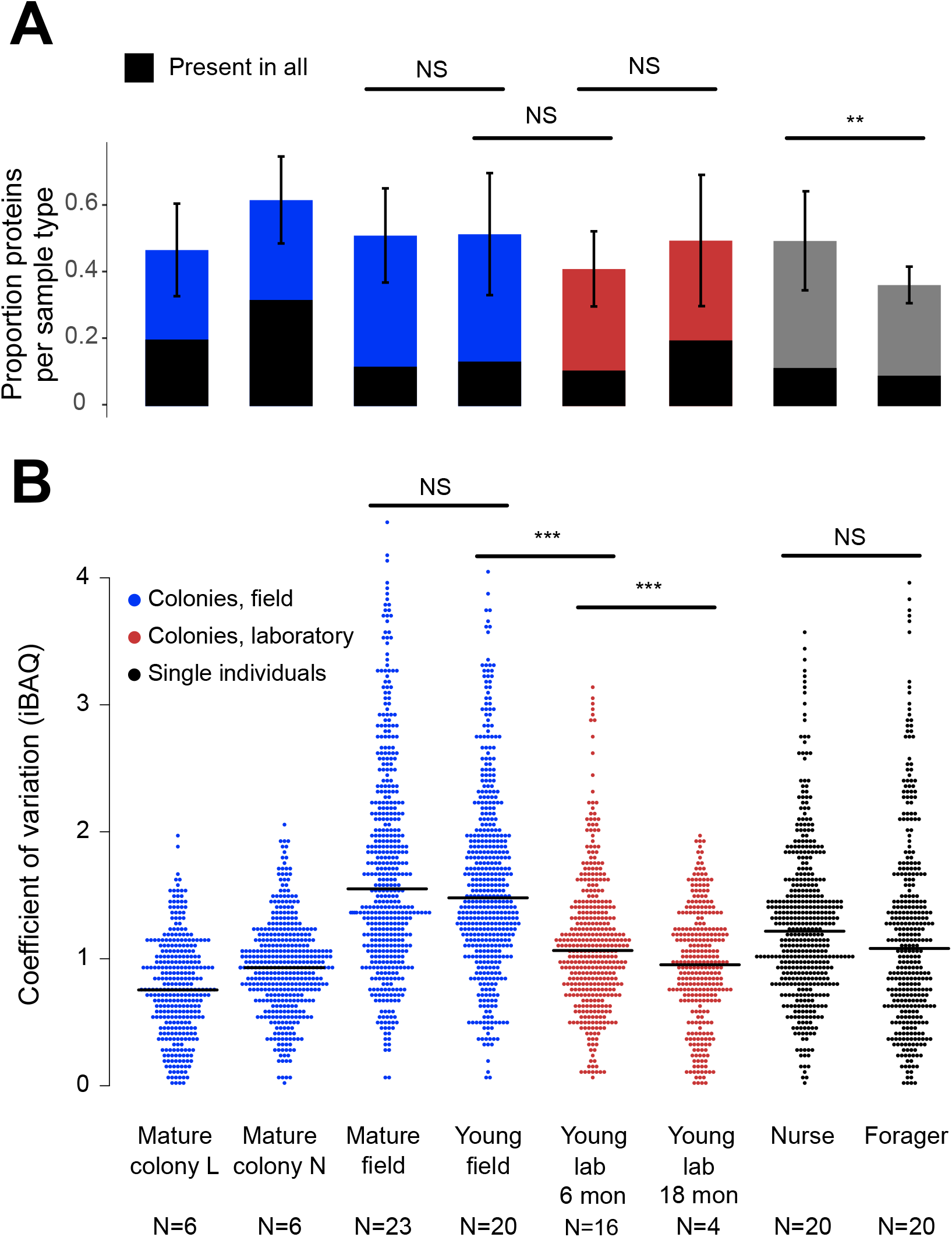
Protein presence in trophallactic fluid varies with biotic and abiotic factors. A. Mean ± SD of the proportion of proteins present in samples of a given type. Proportion of proteins present in all samples of a given type are highlighted in black. B. Coefficient of variation (standard deviation/mean), calculated for the iBAQ values greater than zero of all the proteins identified by sample type. Sample sizes per type are given under their names. Mature L and Mature N are mature colonies that were sampled six times to assess within-colony variation in colony samples. Significance of comparisons based on gamma GLM (A) or negative binomial GLM (B): NS indicated when p > 0.05 significant, ** p <0.01, *** p < 0.001 (full results in SI Table 2).

Field-collected samples exhibited more variable proteomes than did lab-collected samples (Figure 2b, gamma GLM posthoc p-values < 0.001, SI Table 2). Further, colonies that had been in the lab for more than one year showed less variable proteomes than did colonies that had been in the lab for only six months (Figure 2b, gamma GLM z=-4.46, SE=0.04, p <0.001). The trophallactic proteome variability of young and mature colonies did not differ significantly, nor did nurses’ and foragers’ (Figure 2b). Foragers had fewer identified proteins in their trophallactic fluid than did nurses (Figure 2a, negative binomial z=3.72 SE=0.08 P=0.005), and there were no significant differences among the main full colony samples (SI Table 3).

When the principal components of the trophallactic fluid proteomes were analyzed, the samples tended broadly to align with others of the same type, though clusters were not fully distinct (Figure 3A and 3B). We developed a metric, self-similarity (S), to assess the depth of difference within and across sample types (Figure 3C). Because field-collected samples had more diverse protein content even within sample types (Figure 2, 3A), the self-similarity in the *Young vs. Mature* comparison is low (Figure 3C). Single individual samples, and especially forager samples were less complex, allowing a larger proportion of their dissimilarity to be explained by sample type. Further, because our classification of nurse and forager is based on the individuals’ location on brood or out-of-nest, it is possible that some nurse-classified individuals were either misclassified or had trophallactic fluid in their crop uncharacteristic of their behavioral caste.

**Figure 3:**
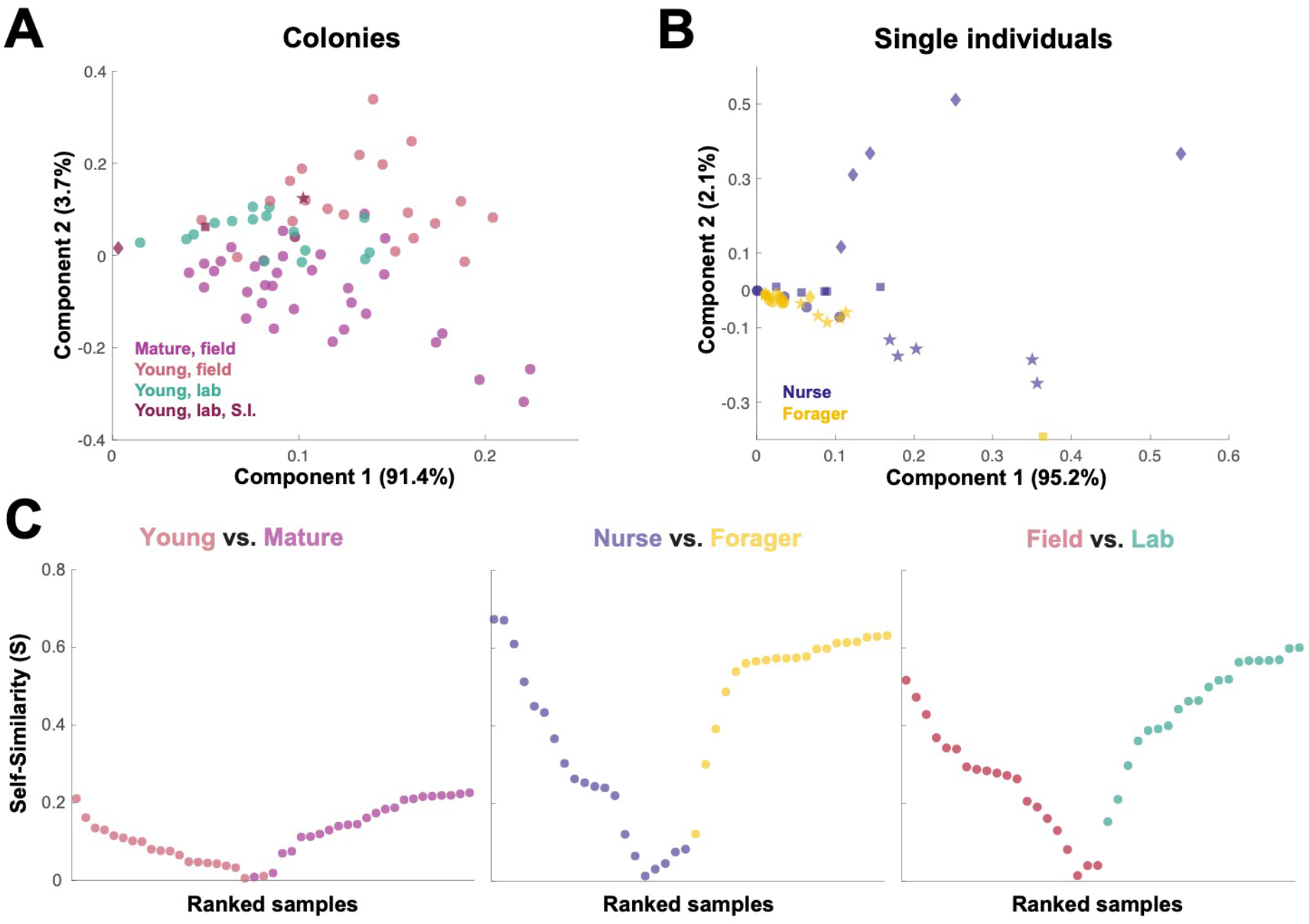
Similarity across trophallactic fluid proteome samples of colonies and single individuals. Principal component analysis for all proteins for (A) colony samples and (B) single individual samples from the four colonies. Symbols representing the four colonies represented in (B) can be found in maroon in (A). C. Ranked Self-similarity S for each sample type comparison. Self-similarity is the absolute value of the difference between dissimilarity within and across samples divided by the average dissimilarity of all samples (by standardized Euclidean distance of protein abundance). Samples with higher S are more similar to samples of the same type, while samples with an S of zero are equidistant to the centroids of the two sample groups.

### Comparisons of trophallactic fluid across conditions

In addition to characterizing the most abundant and core proteins of the trophallactic fluid (Figure 4), we wanted to robustly identify proteins that differ significantly in our comparisons despite the noise inherently present in this social fluid. To accomplish this, we chose to overlay three distinct statistical approaches (Figure 1B): classical frequentist, empirical Bayes and machine-learning in the form of random forest classification. In our main comparisons, *Young vs. Mature* colonies from the field, young colonies in the *Field vs. Lab*, and individual *Nurses vs. Foragers* in the lab, we found significant differences between groups with all three analysis methods (Figure 5, SI Figure 2, SI Figure 4, full results for the significantly differing proteins in Supplemental File 1 and for all proteins in Supplemental Files 2-4).

**Figure 4 (on the following page):**
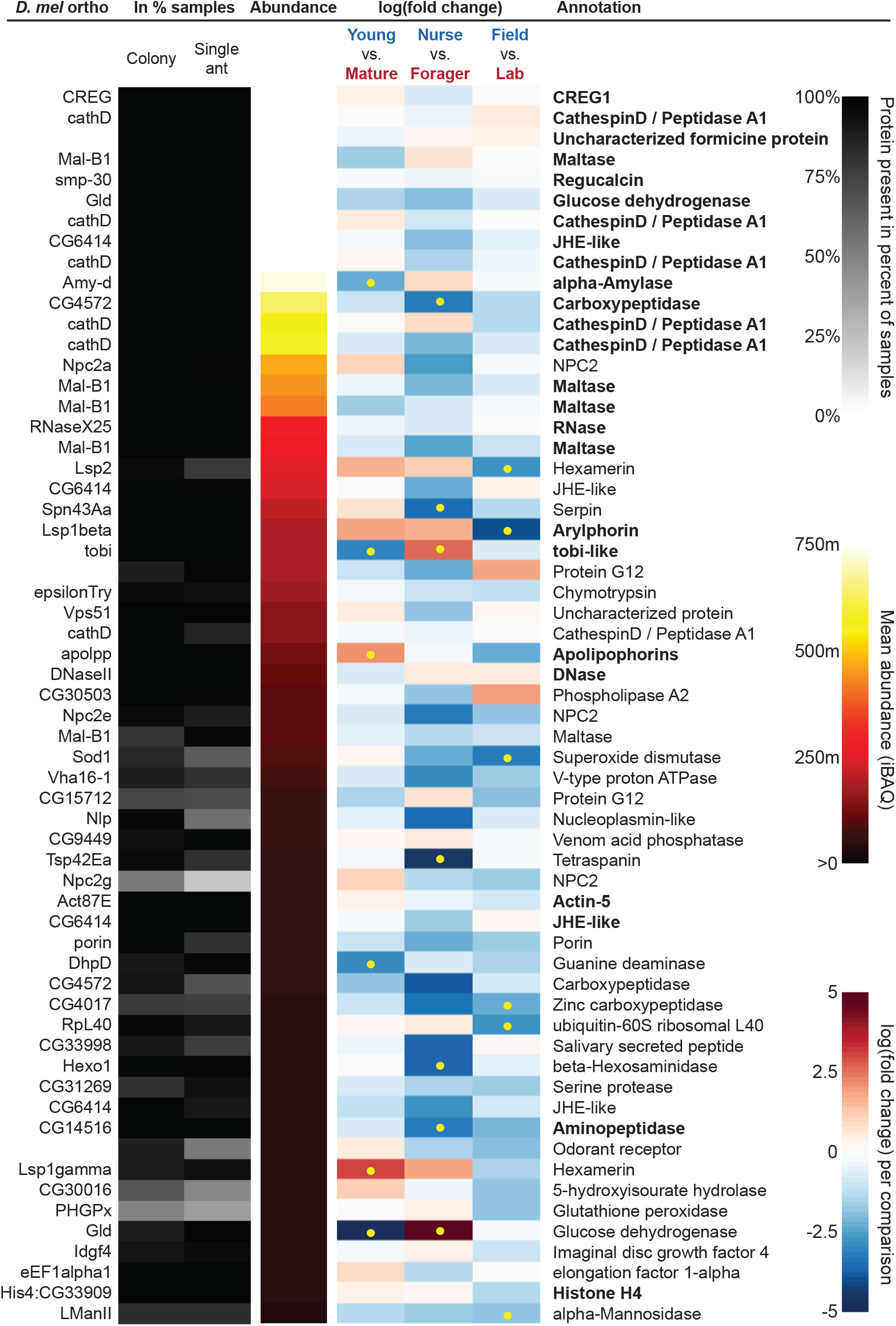
The sixty most abundant proteins in trophallactic fluid over 73 colony and 40 single individual samples. Ranking of abundance (including missing values). From left to right, *Drosophila melanogaster* orthologs, proportion of samples in which the protein was identified in colony samples and single individual samples, average iBAQ abundance across all samples, log2 of the fold change in abundance between types for a given comparison, the comparisons for which the protein was significant in two out of three methods are marked with yellow dots, annotation terms. Annotation terms are bolded for the 25 out of 27 core trophallactic fluid proteins that are amongst the 60 most abundant proteins. The additional but less abundant core proteins are a cathepsin (26-29-p) and a myosin heavy chain (Mhc). For protein accession numbers, see SI Figure 3.

**Figure 5:**
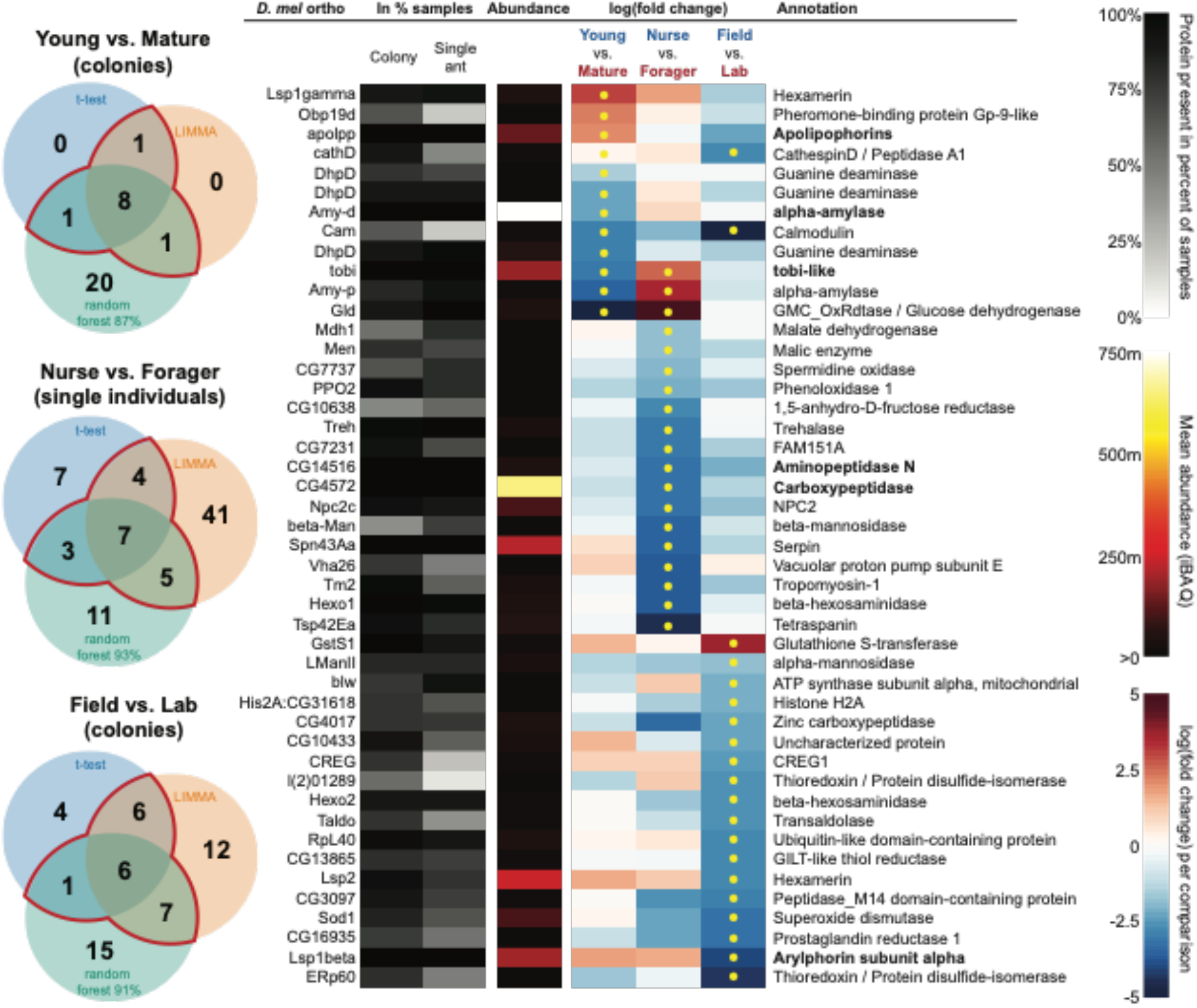
All proteins that significantly differ in two out of three of the analysis methods (frequentist, empirical Bayes and random forest classification with SHAP values). From left to right, Venn diagrams of significance overlap between methods, *Drosophila melanogaster* orthologs, proportion of samples in which the protein was identified in colony samples and single individual samples, average iBAQ abundance across all samples calculated without missing values, log2 of the fold change in abundance between types for a given comparison, the comparisons for which the protein was significant in two out of three methods are marked with yellow dots, annotation terms. Annotation terms are in bold for the core trophallactic fluid proteins present in all samples. For protein accession numbers, see SI Figure 4. For visualization of each analysis method, see SI Figure 2. For all the 135 proteins significantly differing in any analysis, see Supplemental File 1. For full model results, see Supplemental Files 2-4.

For the *Young vs. Mature* comparisons, there were 10, 10 and 30 differentially abundant proteins according to frequentist t-test, empirical Bayesian LIMMA and the random forest approach, respectively. Similarly, for the *Nurse vs. Forager* comparison there were 21, 57 and 26 differentially abundant proteins, and when young colonies were brought to the laboratory and resampled after six months, 17, 31 or 29 proteins had significantly different abundance. The average accuracies of classification for comparisons with the random forest approach were: *Young vs. Mature*, 87%; *Nurse vs. Forager*, 93%; and *Field vs. Lab*, 91%. We found no clear signature of spatial structure (*East vs. West*) in the trophallactic fluid proteomes. The frequentist analysis between different sampling areas found no significantly different proteins, and the random forest model did not reach high enough accuracy for this dataset to be informative (58% classification accuracy). Only the empirical Bayes approach found eight proteins that significantly differed between the sampling areas (SI Figure 2, Supplemental File 1 and 2).

To leverage the unique benefits of the different forms of analysis we focused our further analyses on proteins significantly different in two out of the three forms of analysis. Here, young and mature colonies differed by 12 proteins, and nurses and foragers differed by 19 proteins (Figure 5). When young colonies were brought to the laboratory and resampled six months later, the trophallactic fluid proteomes differed significantly by 20 proteins. Additionally, the single individual dataset showed that proteomes are affected both by colony of origin and by behavioral role of the individual, with 60 proteins showing significant interaction between the two factors (Supplemental File 2).

### Functions of the proteins in trophallactic fluid

To investigate the functions of the proteins found in trophallactic fluid, we performed functional enrichment analysis of gene ontology terms, pathways and protein-protein interaction (PPI) networks of the trophallactic fluid proteins’ *Drosophila melanogaster* orthologs. The sixty most abundant proteins in trophallactic fluid (Figure 4) are predominantly involved in the biological processes of carbohydrate metabolism, lipid and sterol transport (Figure 6, SI Tables 4-6, FDR < 0.00038, FDR < 0.0013 and FDR < 0.0087 respectively). The larval serum protein complex was represented by 3 out of 4 members in both the most abundant proteins and in the significantly differing proteins (hexamerins/arylphorins: Lsp1beta, Lsp1gamma and Lsp2). A strong representation of the innate immune system (Reactome pathway FDR < 6.57e-5) was evident as were lysosomal processes (KEGG pathway, FDR < 3.21e-9).

**Figure 6:**
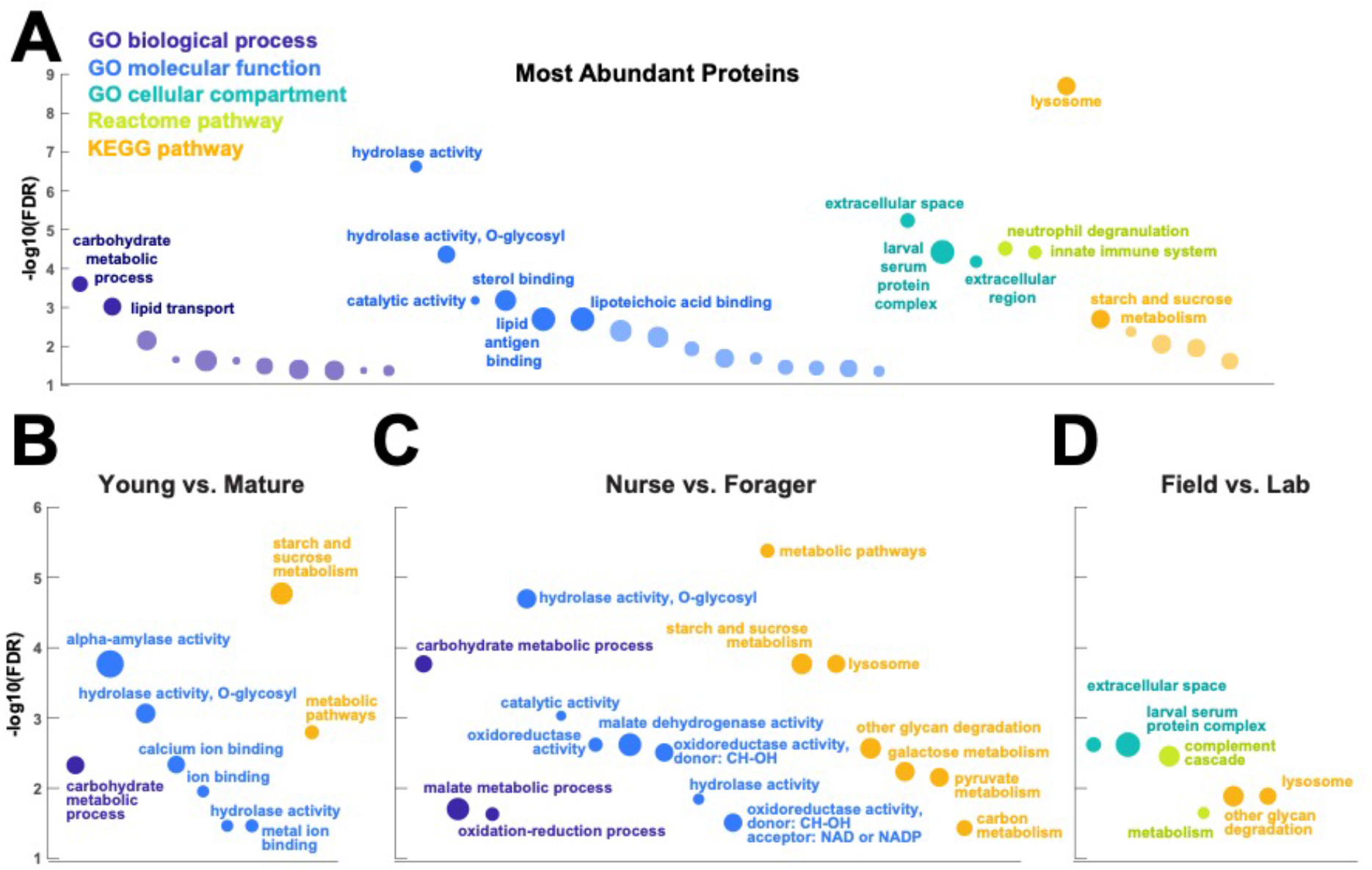
Gene set enrichment analysis of trophallactic fluid. Significant terms for *Drosophila melanogaster* orthologs of (A) the 60 most abundant trophallactic fluid proteins, trophallactic fluid proteins significantly differing between (B) *Young vs. Mature*, (C) *Nurse vs. Forager*, and (D) *Field vs. Lab*, with -log_10_(FDR) indicated on y-axes. Deep purple indicates GO biological process; blue, GO molecular function; turquoise, GO cellular compartment; lime green, Reactome pathway; orange, KEGG pathway. Circle size indicates strength, log_10_(observed proteins / expected proteins in a random network of this size). Full results can be found in SI Tables X-Y.

Beyond the sixty most abundant proteins in trophallactic fluid, many others are of interest as well. A critical protein in insect physiology, vitellogenin is the 93^rd^ most abundant protein in trophallactic fluid, present in 77% and 88% of colony and single individual samples, respectively. Three of the 60 most abundant proteins had no similarity to *Drosophila* genes, and thus could not be included in the functional enrichment analysis. One of them is a putative odorant receptor, another a G-protein alpha subunit, and the third showed no orthology to characterized proteins. None of these proteins significantly differed in more than one analysis for a given comparison.

Many of the trophallactic fluid proteins, abundant or significantly differing, were represented in trophallactic fluid by multiple genes from the same protein family, in some cases part of tandem repeats in the genome, indicative of relatively recent evolution. Multiple proteins of the same family were found in the most abundant trophallactic fluid proteins (Figure 4): a family of cathepsinD-like proteins (six in the top 60; (*18*, *32*) and a family of Maltase-B1-like proteins (five in the top 60). In the list of significantly differing proteins (Figure 5), we observed fewer members of these families and instead saw three guanine deaminase proteins, all of which significantly differed in the *Young vs. Mature* comparison. Other families that showed duplications were glucose dehydrogenases, CREG1 and tobi-like proteins (target-of-brain-insulin).

There was an overlap of 16 proteins between the most abundant proteins and the proteins significant in two out of three of our statistical methods in any of the comparisons. The PPI network for our differentially abundant protein set (46 proteins, Figure 5) was similar to that of the most abundant proteins (Figure 4) but with increased interaction in the networks of the proteins themselves beyond what would be expected by chance (PPI enrichment p-value < 2.35e-11 in differentially abundant proteins relative to p-value < 1.59e-9 in abundant proteins), with noted enrichment in oxidation-reduction processes (FDR < 0.0026) and stronger enrichment in carbohydrate metabolic processes (FDR < 2.15e-6).

To better understand the functions of the significantly differing proteins in each comparison, we analyzed the GO terms and PPI networks of proteins significant in two out of three statistical methods separately for each of our three main comparisons (Figure 6). The *Nurse vs. Forager* comparison yielded a network of proteins with more interaction than would have been predicted by chance (PPI enrichment p-value < 2.57e-4) as well as a higher degree of PPI enrichment than the other two comparisons (Young v Mature p < 0.002 and Field v Lab p < 0.02). The orthologs of differentially abundant proteins found in the behavioral caste comparison involved not only carbohydrate processing (FDR < 1.7e-4), but also oxidation-reduction and malate metabolic processes (FDR < 0.023 and FDR < 0.02, respectively). These pathways have been implicated in the determination of lifespan (*33*). Indeed, two of the 46 differentially abundant proteins over all comparisons have *D. melanogaster* orthologs with the gene ontology term ‘determination of adult lifespan’ (Men, Sod1). The *C. floridanus* tetraspanin, significantly more abundant in nurse trophallactic fluid, is a one-to-many ortholog to the family of Tsp42E genes, one of which has also been implicated in determination of adult lifespan in *D. melanogaster*.

As trophallactic fluid samples of young and mature colonies were distinguishable by principal component analysis and our random forest classifier, we wanted to see if our trained classifier could assess a change in maturity of our young colonies after they had spent six months in the laboratory. Our random forest classifier assigned an average out-of-box maturity score to our 16 laboratory samples of 42% mature, reflecting the intermediate position of the laboratory colony samples in Figure 3.

## Discussion

When an ant colony matures, the protein content of trophallactic fluid changes, suggesting that these proteins that circulate amongst individuals play fundamental roles in age-related colony metabolism and physiology. In ant trophallactic fluid, there are core proteins that are always present but can change in abundance, and proteins whose presence varies based on colony life cycle and environmental context. At the individual level, certain proteins correlate with behavioral caste within the colony, a trait known to encompass both individual task requirements and age (*34*–*36*). Certain overarching patterns across our comparisons are indicative of function: Proteins abundant in mature colonies and nurses are likely factors required to build large workers and sexuals. Proteins abundant in foragers and young colonies are likely to be molecular machinery for fast nutrient uptake. The trophallactic fluid proteomes reveal a rich network of proteins involved in the principal metabolic functions of ant colonies and their life cycle.

The trophallactic fluid complexity declines over time when colonies are brought from the field to the laboratory, showing that the fluid reflects environmental complexity – typical of traits that have evolved to deal with specific environmental cues and stressors (e.g. immunity, (*37*). In our comparisons between young and mature colonies, and between the behavioral castes of individuals within colonies, the overall level of variation stays the same. However, many specific proteins differ in abundance in these comparisons, reflecting adaptation and the functional role of these proteins in the social physiology of ant colonies.

Considering the protein machinery transmitted over the social circulatory system over different temporal and spatial scales reveals fundamental priorities of social insect colonies: controlling the development and life cycle of the whole colony, controlling how individuals work with resources, and finally how evolution has acted as this social fluid has evolved.

### Priorities for resource use as colonies mature

Maximizing resource usage is fundamental for maximizing fitness, and thus is directly linked to life-history evolution and ageing (*33*, *38*, *39*). Our results link trophallactic fluid to the control of the colony life cycle through significantly differing proteins involved in resource use and storage between young and mature ant colonies. The main axis separating young and mature colonies is the tradeoff between maintenance and reproduction (Gadgil & Bossert, 1970): young colonies focus on rapid growth and maintenance, whereas mature colonies consolidate resources into more resource-expensive individuals, sexuals, i.e. for reproduction and dispersal.

Many of the proteins that significantly differ in trophallactic fluid between young and mature colonies correspond to those priorities – resource transmission and consolidation across individuals through nutrient storage proteins like hexamerins and apolipophorins in mature colonies (*40*–*42*), and rapid usage of and response to nutrients through tobi, glucose dehydrogenase, and amylases in young colonies (*43*, *44*). While the observed difference could partially reflect different environmental conditions and resources used by young and mature colonies, these findings suggest that trophallaxis and the social circulatory system are tools to share resources, share information and make collective decisions on whether to produce more workers or start producing sexuals, tilting the colony from one life-stage to another.

### Individual ageing and soma-germline balance

Viewing the colony as a superorganism, the division of reproductive labor between soma and germline should result in different individuals requiring differing resources and sustaining differing metabolic costs. Trophallactic fluid contains numerous factors linked to ageing and coping with oxidative stress, including two of the three most well-known antioxidant enzymes: superoxide dismutase and glutathione peroxidase (*45*). Considering these and other proteins and pathways that have been implicated in aging, like redox pathways and malate metabolism (*33*, *46*), nurses (closest to the queen in the trophallactic network) show the strongest relative abundance of anti-aging proteins. This links trophallactic fluid to one of the main topics of evolutionary ecology: the longevity-fecundity tradeoff between the metabolic needs for reproduction and coping with consequent oxidative stress (*45*, *47*, *48*). Social insect individuals seemingly escape this tradeoff with long-lived and highly reproductive queens and short-lived, non-reproductive workers (*45*, *48*, *49*), but viewing the colony as the organism reveals instead a possible distributed metabolism.

Our results show that both metabolic and oxidative-stress related factors can be shared over the social circulatory system, potentially playing a central role in explaining why social insects seem to subvert this tradeoff. If molecules dealing with oxidative stress can be spread over the circulatory system, certain individuals may bear the costs that others in the network incur. This could potentially account for some of the puzzling results of previous studies on senescence in social insects, especially in relation to its apparent plasticity (*50*–*52*), and provides a new perspective to analyze the regulatory changes of social insect reproductive castes with regard to aging (*34*, *53*–*56*).

The longevity-fecundity trade-off and caste determination in social insects have long been recognized to implicate juvenile hormone, vitellogenin, and insulin signaling (*34*, *39*, *57*, *58*). Yet, beyond the somewhat controversial link between MRJP1/royalactin and EGFR (*29*, *30*), no mechanism has been proposed to enact the robust regulation of colony maturation observed across social insect species. Here we show that the exchange of materials over the social circulatory system can provide that mechanism. This socially transmitted fluid contains nutrient storage proteins, hormones, major regulators of metabolism and major players in pathways involved in aging. While most previous work has focused almost exclusively on gene expression, we show that for species that engage in trophallaxis, expression studies are necessary but insufficient to understand where in the colony the relevant genes are having their effects. It is conceivable that proteins produced in certain individuals could be sequestered by other individuals, allowing transmission of critical materials across superorganismal tissues (*17*). In this view, molecules that exert effects in different individuals than the individuals that have produced them can be considered superorganismal hormones.

### Modulation and transmission of resources within colony

Ant trophallactic fluid includes proteins dealing with metabolism and nutrient usage and storage, notably transmission of large storage proteins (Lsps, apolpp) and rapid usage of sugar resources (Gld, tobi, Amy, Mal). Many of these metabolism-related proteins are core trophallactic fluid proteins present in all samples in high abundance, but many also differ significantly among the colony and individual life stages in our study, showing that optimizing metabolism over levels of organization and life stages can happen through trophallaxis. Similar shifts can be seen in the tissues of multicellular organisms as life-stage priorities change, for example in the midgut of drosophila females after mating, where significant changes in expression are observed in many of the same genes we note here (*44*). Some targets of these proteins in trophallactic fluid may be adult physiology and others larval physiology. The *Drosophila* larval hemolymph proteome changes as development unfolds, in particular around metamorphosis where nutrients need to be mobilized and repurposed (*59*). Many of these significantly differing proteins over larval development (*59*) also appear in our comparisons, suggesting that this regulation may occur over the social network in some ants. Metamorphosis involves substantial autophagy and autophagic flux, repurposing nutrients through lysosomal pathways (*60*) that are well represented in our abundant and significantly differing protein lists as well.

The protein composition of trophallactic fluid, including many of the above-mentioned metabolism and redox proteins, differs between the nurses and foragers, suggesting that different behavioral castes have different roles at the molecular level as well. We confirm that trophallactic fluid indeed plays a role in the social physiology of the colonies, with different functions distributed among the individuals, similarly to multicellular organisms distributing functions among organs (*17*, *61*). Overall, most of the significantly different proteins have higher abundances in nurses, who also have overall a greater diversity of proteins in trophallactic fluid than do foragers, likely reflecting their role in larval care and development. An earlier study found that in honey bees, foragers have greater expression of immune response, sugar processing and detoxification genes especially in glands in and around the head, tissues dealing with nectar transfer and trophallaxis (*62*). Many of the genes differentially expressed in foragers in their study were more abundant in trophallactic fluid of foragers in our study (namely sugar processing enzymes and glutathione-S-transferases), and our analysis of Reactome pathways in the abundant trophallactic fluid proteins showed significant enrichment in innate immunity genes.

### Evolution of social fluids

Trophallactic fluid is one of many social fluids in biology – milk and seminal fluid are similar examples of direct transfers of biological material between individuals. Such socially exchanged materials often contain molecules that target receivers’ physiology beyond the fundamental reason for the transfer (*63*, *64*). Some of the proteins we find to be important in our comparisons have previously been implicated in these other social transfers. For example, one of our protein hits is orthologous to Drosophila’s CG10433, a seminal fluid protein (*65*) that impacts juvenile-hormone-associated hatch-rate post-mating (*66*). An interesting aspect of ant societies is that the males are generally not present in them, except as genes functioning in workers. Thus it seems possible that some genes traditionally considered to be only male-expressed (such as those coding for seminal fluid proteins like CG10433 (*65*)) could have evolved to have new, paternally-imprinted functions in workers, as a way for the male genome to impact the society life (*67*, *68*).

In another parallel to a phylogenetically distant social fluid, trophallactic fluid’s most abundant protein is a CREG1, a secreted glycoprotein also abundant in mammalian milk (*69*). Another CREG1 protein is significantly more abundant in our field colonies. Additionally, the connection to the previously mentioned gene expression changes in *Drosophila* midgut after mating (*44*) suggests that genes involved in a physiological response to one social fluid (sperm) may have evolved to be important also in connection to another (trophallactic fluid). Beyond these social fluids, there are also exceptional parallels to venom, a fluid evolved for interspecific interaction (*70*, *71*). These kinds of similarities between social fluids may indicate that certain molecular machinery is required to build functional transmitted fluids. Finding molecular parallels in distinct behavioral processes begs for further investigation on the role of such exchanges in the evolution of social physiology. If individuals affect each other’s behavioral trajectories and development in ways that have fitness relevance, this substantially complicates the pathways of inheritance, and allow social effects to directly influence the evolutionary process, as indirect genetic effects (*61*, *72*, *73*).

While many of the proteins in trophallactic fluid are secreted extracellular proteins whose orthologs are also clearly secreted extracellular proteins (e.g. maltases, amylases, hexamerins, superoxide dismutase), many others are classically considered to function within cells either in cytosol, in membranes or specific organelles (e.g. cathepsins, ATP synthase, porin, tetraspanin). Many of these not-classically-secreted proteins are critical players in the secretion machinery, especially in high secretion tissues (*74*). Consistent with roles in secretion we also find markers of exosomes, extracellular vesicles typically present in social fluids (e.g. tetraspanin (*76*)). The presence of canonical and non-canonical markers of secretion indicate that secretion is fundamental for the function this social fluid.

Lysosomal pathways are enriched in our most abundant trophallactic fluid proteins and in our set of significantly varying trophallactic fluid proteins between nurses and foragers according to the KEGG analysis. Lysosomes are clearly acidic and can be major players in secretion, autophagic flux and exocytosis (*77*–*79*) – processes that may be important for nurses that feed larvae by trophallaxis. In general, the lysosomal signatures we see in trophallactic fluid may be traces of the mechanism of secretion, or perhaps be evidence of how this fluid has evolved. As trophallactic fluid has become acidified in formicine ants (*80*), lysosomal genes may have been duplicated and neofunctionalized to a new role in this acidic fluid. This would be a similar repurposing to what was observed with the multiple juvenile-hormone-esterase-like proteins in trophallactic fluid (*19*). The fact that many abundant trophallactic fluid proteins represent clusters of related proteins from a few families (cathepsins, guanine deaminases, maltases) suggest there has been recent evolution in the proteins arriving in this fluid. This hypothesis can be tested by analyzing foregut fluids over the ant phylogeny, between species who engage in trophallaxis and those that don’t, from different subfamilies including those without an acidified gut. It is an exciting future goal to assess how classical metabolism-related proteins have evolved to function in their new role in inter-individual transmission in trophallactic fluid.

A large proportion of the proteins in trophallactic fluid seems to be inducible – their presence and abundance are triggered by certain environmental, social or developmental contexts. The current study focuses on some of these contexts, but there are still many more that lie outside the scope of this work. For example, based on our data from the four colonies we sampled, individuals of different colonies harbor significantly different sets of proteins in their trophallactic fluid, for reasons we cannot assess with the current data. Ant colonies typically have colony signatures in many traits, and when based on genetic differences, these can extend to larger scale spatial patterns due to the often strong population genetic structures of ants (*81*). However, this is not the case here: the trophallactic fluid proteomes do not differentiate between sampling locations although they do differ among colonies, indicating that colony signatures are not just genetic, but also plastic and possibly functional. Previous screens on population proteomics in other study systems have found differences among populations in similar spatial scales (*82*–*84*). As our individual-level samples were collected from laboratory colonies and thus lack many of the environmental reasons for differentiation (*85*), we suggest the most likely explanations are health status and slight differences in the social context (for example brood composition). Pinpointing all contexts that induce changes in trophallactic fluid, along with the exact functions of the proteins, are important subjects for future work.

## Conclusions

We show that the protein composition of ant trophallactic fluid varies across different external contexts and internal conditions both at the colony and at the individual level, suggesting that the changing trophallactic fluid proteome has key functions in social physiology and life cycle of colonies. By describing the natural variation of trophallactic fluid we have laid the groundwork for future studies on the functions of these proteins in controlling the colony life cycle, senescence and behavior.

## Materials and methods

### Study species

*Camponotus floridanus* is a common species of carpenter ant in the south-eastern USA, and has already been the focus of previous trophallactic fluid analyses (*18*, *19*). They live in dead wood or in man-made structures, often in urban habitats, and forage for honeydew, floral nectar, extra-floral nectar, and arthropod prey. Each colony has a single, singly mated queen (*86*), and polydomous nest-structures where queenless satellite nests are common. Colonies grow to tens of thousands of workers and produce sexual brood only after multiple years of initial growth. Large established colonies have two morphologically differentiated worker castes, with variably sized small-headed minors focusing on brood care when young and foraging when old, and big-headed majors that engage in nest defense, foraging and food storage (*87*).

### Colony and sample identification

The species was identified based on worker and queen morphology (*87*–*89*). In line with previous studies, we use the name *C. floridanus* with the knowledge that the taxonomy and nomenclature of the C. *atriceps* complex (to which it belongs) is not fully resolved (*87*).

We collected full young colonies (0-80 workers) and mature colony extracts (30-200 workers) on several Florida Keys islands (Figure 1 and SI Table 1) in winter 2019 and 2020. A colony was deemed “young” if the worker population was <100, primarily minors, and the queen was found (meaning both that the species could be clearly identified and that the nest was not a queenless satellite of an established colony), and “mature” if the colony was larger (>1000 individuals visible) and the opened nest contained many large aggressive majors. Young colonies lack majors (*90*) and individuals are generally less aggressive. We only collected mature colony samples when we also found larval brood in the opened nest. In our study area, we observed that young colonies are typically found nesting in different material than are mature colonies. Young colonies are often found under stones or in lumps of clay-like mud associated with crab burrows a short distance from the water, whereas the mature colonies were found nesting in large pieces of damp rotting wood.

### Laboratory rearing

Young colonies were brought to the lab and maintained in fluon-coated plastic boxes with a mesh-ventilated lid, at 25 °C with 60 % relative humidity and a 12 h light/dark cycle. Each colony was provided with one or more glass tube for nesting, 10% sugar water, and a Bhatkar & Whitcomb diet (*91*) and some *Drosophila melanogaster*. One week prior to proteomic sampling, we substituted the honey-based food with maple syrup-based food to avoid contamination with honeybee proteins (as in LeBoeuf et al 2016).

### Trophallactic fluid collection

Field samples of trophallactic fluid were collected within eight hours of ant collection. Of the 20 young colonies and 23 mature colonies, workers collected from two of the mature colonies (L and N) were subdivided into six fragments to assess variation within a single colony (two samples from major workers, two samples from brood-associated workers, and two samples from the remaining minor workers). For all other analyses only one of these for each colony (referred to as minors1) was used to avoid pseudo-replication. In the laboratory, the trophallactic fluid samples underlying the *Field vs. Lab* comparison were sampled after six months in the lab. The four colonies used for the single individual analyses had been in the lab for 18 months at the time of trophallactic fluid collection.

Trophallactic fluid was obtained from CO_2_- or cold-anesthetized workers whose abdomens were gently squeezed to force them to regurgitate the contents of their crops (as in *18*). For each colony, at least 30 individuals were sampled to obtain at least 10 μl of raw trophallactic fluid. For many young colonies only smaller samples were possible, because of the low number of workers (SI Table 1). Young colony samples were only used for further analysis if at least 2.5 μl of trophallactic fluid were collected. For single individual samples, workers with visibly full abdomens were chosen and the obtained sample volumes ranged from 0.7 μl to 2.2 μl. An individual was classified as forager, when it was seen outside the nest tube in the feeding area of an undisturbed laboratory nest box, and a nurse, when it remained in the nest tube even after the tube was removed from the original laboratory nest and placed into a new one. For colonies from which individual samples were collected, a pooled sample was also taken from individuals that remained after individual sampling. Samples were collected with glass capillaries into 5 μl of 1 x Sigmafast Protease Inhibitor Cocktail (Sigma-Aldrich) with 50 mM Tris pH 9 in LoBind eppendorf tubes and were stored -80 C until further analysis. The total proteomics sample number is 73 colony samples of following types: 23 mature colonies with two of them sampled 6 times, 20 young colonies in the field, 16 young colonies in the laboratory, four laboratory colonies used for single individual sampling; and 40 individual samples: 20 nurses and 20 foragers.

### Protein mass spectrometry sample preparation and analysis

Samples were mixed with Laemmli sample buffer and pH was adjusted with 1 M Tris-Cl, pH 7. After reduction with 1 mM DTT for 10 min at 75°C and alkylation using 5.5 mM iodoacetamide for 10 min at room temperature protein samples were separated on 4-12% gradient gels (ExpressPlus, GeneScript). Each gel lane was cut into small pieces, proteins were in-gel digested with trypsin (Promega) and the resulting peptide mixtures were processed on STAGE tips (*92*, *93*).

LC-MS/MS measurements were performed on a QExactive plus mass spectrometer (Thermo Scientific) coupled to an EasyLC 1000 nanoflow-HPLC. HPLC-column tips (fused silica) with 75 μm inner diameter were self-packed with Reprosil-Pur 120 C18-AQ, 1.9 μm (Dr. Maisch GmbH) to a length of 20 cm. A gradient of A (0.1% formic acid in water) and B (0.1% formic acid in 80% acetonitrile in water) with increasing organic proportion was used for peptide separation (loading of sample with 0% B; separation ramp: from 5-30% B within 85 min). The flow rate was 250 nl/min and for sample application 650 nl/min. The mass spectrometer was operated in the data-dependent mode and switched automatically between MS (max. of 1×10^6^ ions) and MS/MS. Each MS scan was followed by a maximum of ten MS/MS scans using normalized collision energy of 25% and a target value of 1000. Parent ions with a charge state form z = 1 and unassigned charge states were excluded from fragmentation. The mass range for MS was m/z = 370-1750. The resolution for MS was set to 70,000 and for MS/MS to 17,500. MS parameters were as follows: spray voltage 2.3 kV; no sheath and auxiliary gas flow; ion-transfer tube temperature 250°C.

The MS raw data files were uploaded into MaxQuant software (*94*), version 1.6.2.10, for peak detection, generation of peak lists of mass error corrected peptides, and for database searches. MaxQuant was set up to search both the UniProt (RRID:SCR_002380, www.uniprot.org) and NCBI (RRID:SCR_003496, www.ncbi.nlm.nih.gov) databases restricted to *C. floridanus* (UniProt, February 2020 version; NCBI RefSeq, version 7.5), along with common contaminants, such as keratins and enzymes used for digestion. Carbamidomethylcysteine was set as fixed modification and protein amino-terminal acetylation and oxidation of methionine were set as variable modifications. Three missed cleavages were allowed, enzyme specificity was trypsin/P, and the MS/MS tolerance was set to 20 ppm. The average mass precision of identified peptides was in general less than 1 ppm after recalibration. Peptide lists were further used by MaxQuant to identify and relatively quantify proteins using the following parameters: peptide and protein false discovery rates, based on a forward-reverse database, were set to 0.01, minimum peptide length was set to 7, minimum number of peptides for identification and quantitation of proteins was set to one which must be unique. The ‘match-between-run’ option (0.7 min) was used, which helps improve the protein identifications especially for our single-individual samples. All proteins labelled as contaminants, reverse or only identified by site were excluded and proteins with scores less than 70 were removed. After the filtering, the dataset contained 519 proteins. Quantitative analysis was performed using iBAQ values. Intensity-based absolute quantification (iBAQ) is the quotient of sum of all identified peptides and the number of theoretically observable peptides of a protein (*95*).

### Statistical analyses

Analyses of dataset characteristics were performed in Perseus v1.6.15.0 (*96*), R 3.6.1 (*97*) and Matlab R2020b (Figures 2 and 3). Differences in protein numbers among the sample types were analyzed with a negative binomial model, using the function nb.glm from the R-package MASS 7.3-53 (*98*). Proteome variability per sample type, as measured by the coefficient of variation of the iBAQ abundance of each protein when present, was analysed with a generalized linear model with gamma distribution and log-link with the R-package LME4 (1.1-26) (*99*). The package multcomp 1.4-15 was used for post-hoc testing for both models. Pearson correlation tests were used to check whether obtained protein number correlates with the sample volume. Because significant correlation was found, all further analyses were done separately for the individual samples that have small volume, and colony samples that have larger volume. Principal component analysis was run in Matlab on raw iBAQ values, for both the individual and the colony datasets.

Metric for self-similarity (S) within and across samples was calculated in Matlab2020b as follows: pairwise standardized Euclidean distances (dissimilarities, D) were calculated between each pair of samples based on square-root transformed and median subtracted protein abundances; these dissimilarities were averaged for each sample with the other samples within type 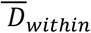 and with the samples of the other type 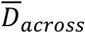 and divided by the average dissimilarity to all other samples. Thus, self-similarity was calculated as:

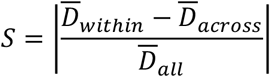

To establish the proteins whose abundance differs significantly between sample types, samples were subdivided according to three main comparisons (Figure 1): *Young vs. Mature* colonies from the field, young colonies in the *Field vs. Lab* six months later, and individual *Nurses vs. Foragers* in the lab. In addition, the extent of spatial effects was analyzed for the field-collected *Young vs. Mature* dataset by dividing the sampling locations to two areas (*East vs. West*). For the colony data, the differing sample volumes may account for a small proportion of the significant differences in the *Young vs. Mature* comparison, and to lesser extent in the *Field vs. Lab* comparison, where sample volume is collinear with the sample type. Our analyses may miss some of the proteins more abundant in the young field collected colonies which have the smallest sample volumes.

Quantitative proteomic comparisons between sample types were performed independently with three different approaches to robustly identify significantly differing proteins: 1) classical frequentist t-tests, 2) linear models with empirical Bayes variance correction, and 3) machine-learning paired with modified Shapley values. Our approach is designed to be at the same time conservative and to find most of the differing proteins among our comparisons of the trophallactic fluid. The frequentist t-tests are the most conservative, and they miss some interesting proteins due to their strict model expectations that allow only to use the most common proteins. The empirical Bayes approach to cope with sample variance is a more flexible method that allows use of the entire dataset, finding important hits also among the rarer proteins, although the high amount of missing values, where iBAQ equals zero, makes the model less powerful for these proteins (*100*). The machine learning approach paired with modified Shapley values, although less well explored in the current proteomics literature, is promising for its ability to find multivariate patterns that the other methods miss, and results in interpretable classification. For each comparison, we report the full results of all three analyses in Supplemental File 1 (significantly differing proteins only) and Supplemental Files 2-4 (all results). Our results and discussion sections focus on the proteins that appear significantly different based on two out of three analysis methods (Figure 5).

### Classical frequentist analysis

Within each dataset only proteins present in over 70% of the samples were analyzed. Out of an original 519 proteins, the final datasets for each comparison contained the following number of proteins: *Young vs. Mature*, 172; *Field vs. Lab*, 137; and *Nurse vs. Forager*, 136. All data were log2 transformed and median-centered, and missing data were imputed by random sampling from normal distribution with 2SD downward shift and 0.3 width for each sample. For colony datasets, we used the permutation-based FDR of 0.05, and for the single individual dataset that contained more borderline-significant proteins, we used a more stable Benjamini-Hochberg FDR with a stricter threshold of 0.01. S0 parameter (similar to fold-change) was set to 2 for all analyses. All comparisons were run as two-sample t-tests, with the *Field vs. Lab* as paired.

For the individual dataset, the combined effects of colony identity and behavioral role (*Nurse vs. Forager*) and their interaction were analyzed with two-way ANOVA, with Benjamini-Hochberg FDR corrections performed in R with the base R 3.6.1 command ‘p.adjust’. Both factors were also analyzed separately with multiple- and two-sample t-tests (for colony identity and behavioral role, respectively). To allow comparison to the other statistical methods, only the simple *Nurse vs. Forager* analysis without the interaction was used for combining the lists of significantly different protein abundances. Our balanced sampling guarantees the results of this simpler model are robust enough to find the most descriptive proteins for nurse and forager trophallactic fluid, even when the more complex interactive patterns are lost.

### Empirical Bayesian analysis

We implemented LIMMA (Linear Models for Microarray Data), a method for two-group comparison using empirical Bayes methods to moderate the standard errors across proteins (*100*), on our score-filtered iBAQ proteomic datasets with the LIMMA-pipeline-proteomics pipeline 3.0.0 (http://doi.org/10.5281/zenodo.4050581) developed for R 4.0.2. Data were median-normalized before comparison and all comparisons were run with a log2 fold change cutoff of 2.

### Random forest and SHAP analysis

We used random forest models (sklearn.ensemble.RandomForestClassifier version 0.22.1, Pedregosa et al., JMLR 12, pp. 2825-2830, 2011) to classify samples into one of two groups for each comparison. These analyses were performed in Python 3.7.6 in a Jupyter notebook. For each comparison, ten analyses were performed, each with a different seed. For each seed, the dataset was split into 80% training set and 20% test set, and a model was fit, tested and accuracy computed. If accuracy was below 85%, hyper-parameter tuning was performed with GridSearchCV (sklearn 0.22.1), and the model re-fit. A seed and its corresponding model were not retained for further analysis if accuracy could not be improved above 75%. Accuracies for *East vs. West* ranged from 33-89% and over 20 seeds, only one could be improved above 75%. The typical parameters: max_depth, 3 or 5; max_features, ‘auto’; min_samples_leaf, 3; min_samples_split, 8 or 12; n_estimators, 100 or 500. Samples were classified with out-of-box scores (Supplemental File 4). The average accuracies of classification for comparisons were: *Young vs. Mature*, 87%; *Nurse vs. Forager*, 93%; *Field vs. Lab*, 91%; *East vs. West*, 58%.

To understand which proteins contributed to the classification, we used SHAP (SHapley Additive exPlanations, shap package v0.37.0 for Python 3), a game theory tool that explains the output of machine learning models (*101*). To analyze the importance of each protein in a given comparison (feature importance), we averaged the absolute value of the Shapley values per protein across the data to derive the feature importance. Then for each protein, we averaged the feature importances over each of the ten seeded models. Proteins that have no impact on the model classification receive a feature importance value of 0. When ranked according to average feature importance, the data had an approximate Pareto distribution with an inflection point typically at feature importance of ~0.15. Thus, because there is no established cutoff for significance in this form of analysis, we chose to include as ‘significant’ in further analyses all proteins with a feature importance of > 0.15 (Supplemental File 4).

For random forest predictions, models trained on the classification between young and mature colonies were used to classify the same young colonies after six months in the laboratory. Out-of-box scores were averaged over five seeded models.

### Orthology, gene ontology, and protein network analyses

Because little functional work has been done in ants, we analyzed gene ontology terms for the *Drosophila* orthologs to our genes of interest. Orthologs to *C. floridanus* trophallactic fluid proteins were determined with OMA (“Orthologous MAtrix” Jan 2020 release (*102*)). If no ortholog was found within OMA for a given gene, the protein sequence was protein BLASTed against *Drosophila melanogaster*. In some cases, no ortholog could be found. Annotations were compiled from NCBI RefSeq and UniProt annotations.

GO analysis was performed using both Flybase (*103*) and STRING v11 (*104*). STRING was also used for protein-protein interaction and pathway analyses, including KEGG and Reactome (SI Table 4-6). The protein-protein interaction enrichment analysis in STRING used a hypergeometric test with Benjamini-Hochberg corrected FDR. Only 43 out of the 60 most abundant proteins had sufficient annotation for use by STRING while 44 of the 46 differentially abundant proteins had sufficient annotation.

## Acknowledgements and funding information

We thank Joanne Reiter for help with field work, Guillaume Kuhn for help with ant maintenance in the laboratory, and members of the Social Fluids lab for their useful comments on the manuscript.

## Author contributions

Conceptualization: ACL, SH

Methodology: ACL, SH, MS

Investigation: ACL, SH, MPM, MS

Visualization: ACL, SH, MS

Supervision: ACL

Writing—original draft: SH, ACL

Writing—review & editing: ACL, SH, MPM, MS

## Data and materials availability

All data is either made available in this submission and raw proteomics data will be uploaded to proteomXchange.

## SUPPLEMENTAL INFORMATION FOR

**SI Figure 1:**
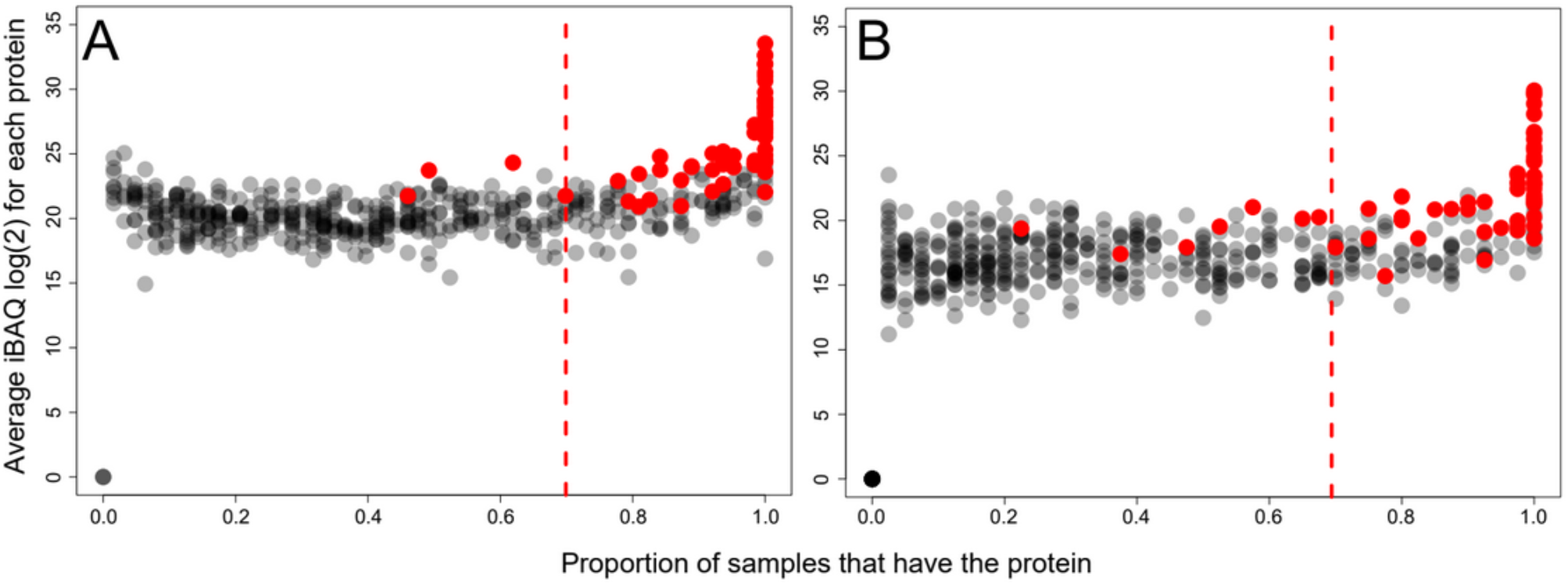
Protein abundance and commonness. Protein abundances of the 519 proteins, calculated without missing values (where no matching spectra were detected), in A) the colony dataset, and B) in the single individual dataset. The proteins highlighted in red are the most abundant ones when calculated including missing values in both datasets combined, as shown in Figure 3. The red dashed line shows the cut-off used for classical frequentist statistical analyses – for the empirical Bayes and machine learning analyses all proteins were included.

**SI Figure 2:**
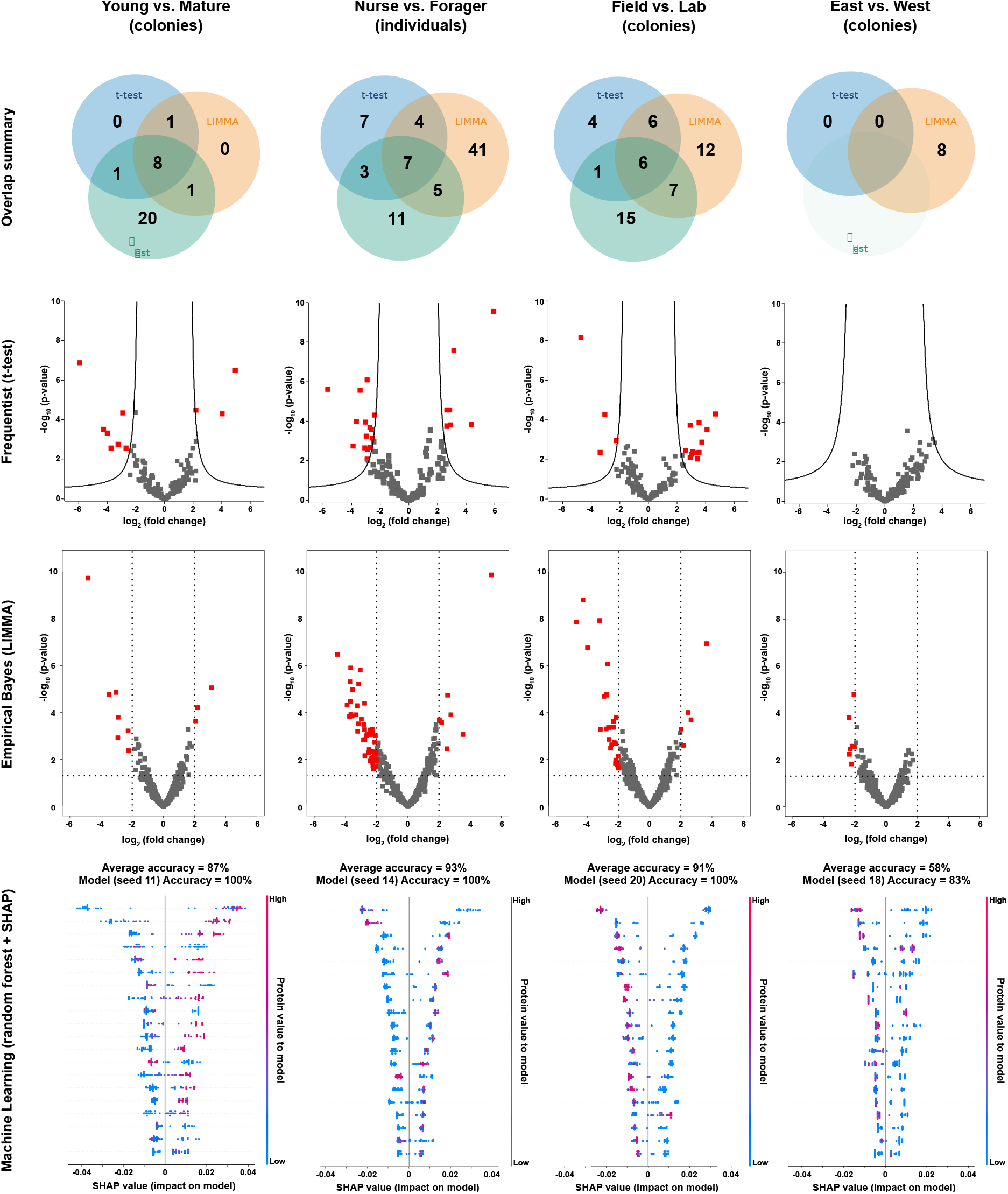
Visualization of all results. Venn diagrams summarizing statistical methods, frequentist volcano plots, empirical Bayes volcano plots, example SHAP value plots of feature importance the top 20 proteins. Each SHAP plot is for one of the ten models trained. For significant proteins, see Supplemental file 1, for full model results, see Supplemental files 2-4.

**SI Figure 3:**
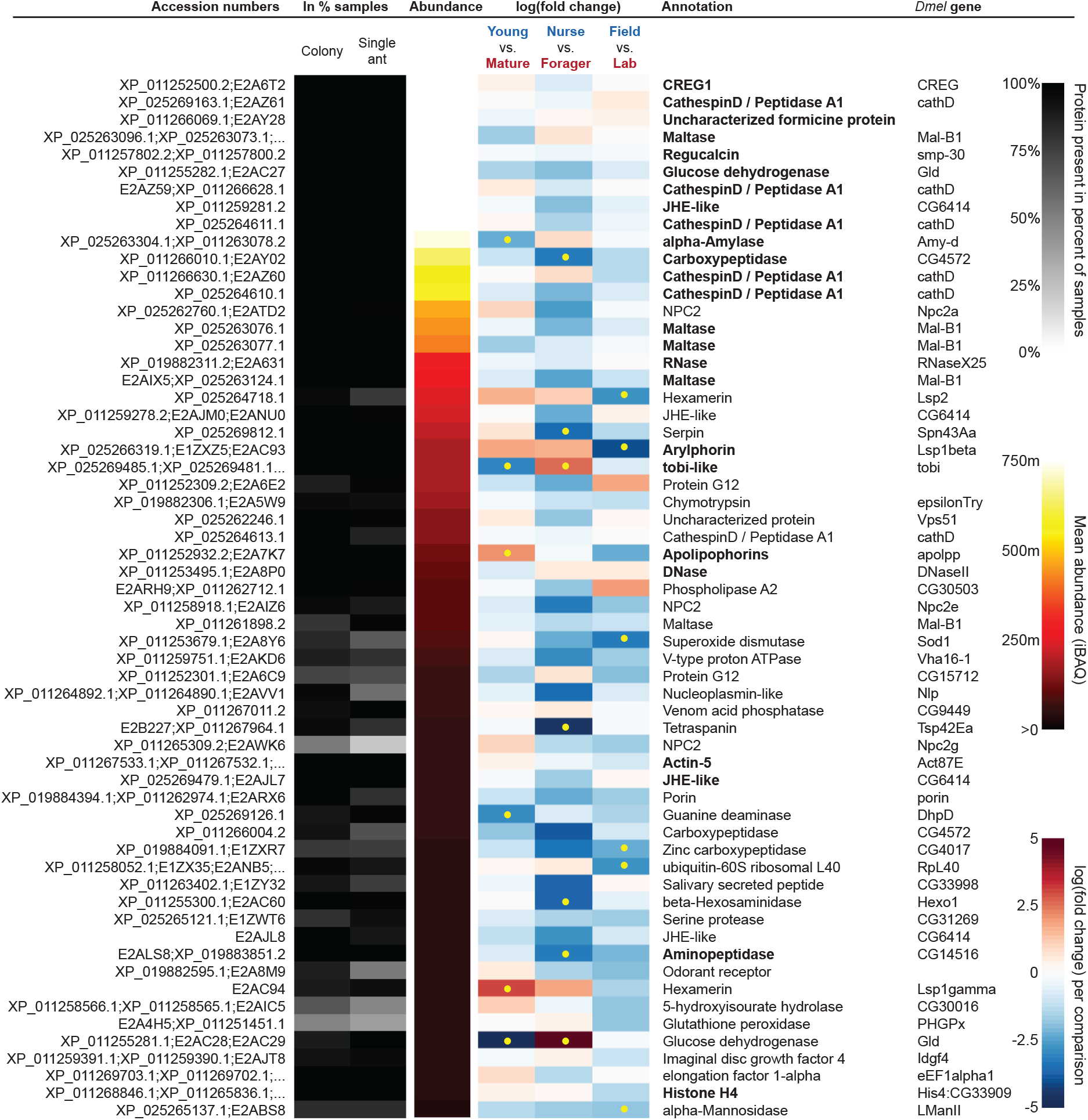
Most abundant proteins with accession numbers. The sixty most abundant proteins in trophallactic fluid over 73 colony and 40 single individual samples. Ranking of abundance included zero values. From left to right, accession numbers, proportion of samples in which the protein was identified in colony samples and single individual samples, average iBAQ abundance across all samples, log2 of the fold change in abundance between types for a given comparison, the comparisons for which the protein was significant in two out of three methods are marked with yellow dots, annotation terms. Annotation terms are bolded for the 25 out of 27 core trophallactic fluid proteins that are amongst the 60 most abundant proteins. The additional but less abundant core proteins are a cathepsin (26-29-p) and a myosin heavy chain (Mhc).

**SI Figure 4:**
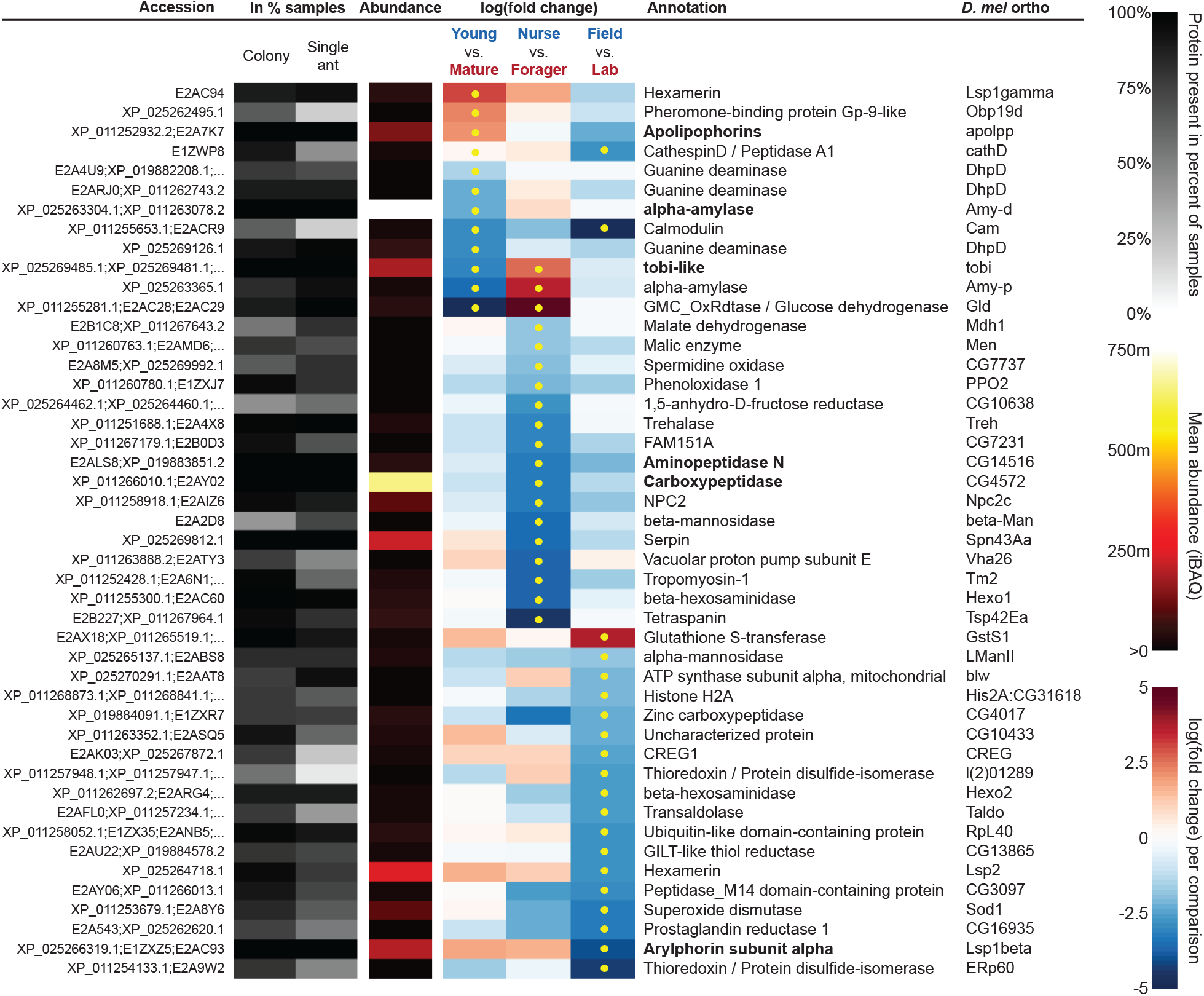
Significantly differing proteins in two out of three analyses with accession numbers. All proteins that significantly differ in two out of three of the analysis methods (frequentist, empirical Bayes and random forest classification with SHAP values). From left to right, accession numbers, proportion of samples in which the protein was identified in colony samples and single individual samples, average iBAQ abundance across all samples calculated without zero values, log2 of the fold change in abundance between types for a given comparison, the comparisons for which the protein was significant in two out of three methods are marked with yellow dots, annotation terms.

**SI Table 1:** The sampling scheme. Trophallactic fluid (TF) sampled for proteomics analysis. Date_field = when the colony extract was collected from the field site, Date_TF = the date of the trophallactic fluid collection. Vol = the volume of trophallactic fluid collected, N (ants) = the number of ants sampled in colony samples. The lab2019 colonies were used for single individual trophallactic fluid samples. For the two mature colonies that were sampled six times, * marks the sample that was used in the main datasets.

| NestID | TFsampleID | Island | Coordinates | Date_field | Date_TF | Type | Vol (μl) | N (ants) |
| --- | --- | --- | --- | --- | --- | --- | --- | --- |
| cflo20.02 | FieldYoung2002 | Fiesta | 24.840091, -80.792447 | Feb 20, 2020 | Feb 20, 2021 | young/field | 5 | 23 |
| cflo20.04 | FieldYoung2004 | Fiesta | 24.840091, -80.792447 | Feb 20, 2020 | Feb 20, 2020 | young/field | 3 | 7 |
| cflo20.07 | FieldYoung2007 | Sugarloaf | 24.627298, -81.569617 | Feb 21, 2020 | Feb 21, 2020 | young/field | 3.3 | 7 |
| cflo20.09 | FieldYoung2009 | Sugarloaf | 24.627298, -81.569617 | Feb 21, 2020 | Feb 21, 2020 | young/field | 2.3 | 6 |
| cflo20.11 | FieldYoung2011 | Tea Table | 24.896367, -80.662087 | Feb 22, 2020 | Feb 22, 2020 | young/field | 2.8 | 6 |
| cflo20.12 | FieldYoung2012 | Conch | 24.786233, -80.896969 | Feb 22, 2020 | Feb 22, 2020 | young/field | 10 | 30 |
| cflo20.14 | FieldYoung2014 | Duck | 24.769611, -80.914853 | Feb 23, 2020 | Feb 23, 2020 | young/field | 6.7 | 18 |
| cflo20.16 | FieldYoung2016 | Sugarloaf | 24.627298, -81.569617 | Feb 24, 2020 | Feb 24, 2020 | young/field | 2.6 | 6 |
| cflo20.18 | FieldYoung2018 | Sugarloaf | 24.627298, -81.569617 | Feb 24, 2020 | Feb 24, 2020 | young/field | 5.3 | 15 |
| cflo20.20 | FieldYoung2020 | Sugarloaf | 24.627298, -81.569617 | Feb 25, 2020 | Feb 25, 2020 | young/field | 10 | 23 |
| cflo20.21 | FieldYoung2021 | Sugarloaf | 24.627298, -81.569617 | Feb 25, 2020 | Feb 25, 2020 | young/field | 6.6 | 12 |
| cflo20.22 | FieldYoung2022 | Sugarloaf | 24.627298, -81.569617 | Feb 25, 2020 | Feb 25, 2020 | young/field | 15 | 29 |
| cflo20.23 | FieldYoung2023 | Sugarloaf | 24.627298, -81.569617 | Feb 25, 2020 | Feb 25, 2020 | young/field | 6.6 | 9 |
| cflo20.26 | FieldYoung2026 | Conch | 24.786233, -80.896969 | Feb 28, 2020 | Feb 28, 2020 | young/field | 15.4 | 22 |
| cflo20.27 | FieldYoung2027 | Conch | 24.786233, -80.896969 | Feb 28, 2020 | Feb 28, 2020 | young/field | 10.2 | 15 |
| cflo20.28 | FieldYoung2028 | Conch | 24.786233, -80.896969 | Feb 28, 2020 | Feb 28, 2020 | young/field | 10 | 9 |
| cflo20.29 | FieldYoung2029 | Conch | 24.786233, -80.896969 | Feb 28, 2020 | Feb 28, 2020 | young/field | 3 | 8 |
| cflo20.30 | FieldYoung2030 | Conch | 24.786233, -80.896969 | Feb 28, 2020 | Feb 28, 2020 | young/field | 10 | 13 |
| cflo20.33 | FieldYoung2033 | Conch | 24.786233, -80.896969 | Feb 28, 2020 | Feb 28, 2020 | young/field | 10.9 | 16 |
| cflo20.36 | FieldYoung2036 | Conch | 24.786233, -80.896969 | Feb 28, 2020 | Feb 28, 2020 | young/field | 12 | 47 |
| A | FieldMatA | Duck | 24.769611, -80.914853 | Feb 19, 2020 | Feb 19, 2020 | mature | >10 | >30 |
| AA | FieldMatAA | Conch | 24.786233, -80.896969 | Feb 27, 2020 | Feb 27, 2020 | mature | 15.8 | 25 |
| BB | FieldMatBB | Conch | 24.786233, -80.896969 | Feb 27, 2020 | Feb 27, 2020 | mature | 19 | 21 |
| F | FieldMatF | Duck | 24.769611, -80.914853 | Feb 19, 2020 | Feb 19, 2020 | mature | 15 | 30 |
| G | FieldMatG | Fiesta | 24.840091, -80.792447 | Feb 20, 2020 | Feb 20, 2020 | mature | >10 | >30 |
| H | FieldMatH | Fiesta | 24.840091, -80.792447 | Feb 20, 2020 | Feb 20, 2020 | mature | NA | 12 |
| I | FieldMatI | Fiesta | 24.840091, -80.792447 | Feb 20, 2020 | Feb 20, 2020 | mature | 16 | 30 |
| K | FieldMatK | Sugarloaf | 24.627298, -81.569617 | Feb 21, 2020 | Feb 21, 2020 | mature | 15 | 30 |
| L_Majors1 | FieldMatLmaj1 | Sugarloaf | 24.627298, -81.569617 | Feb 21, 2020 | Feb 21, 2020 | mature | >10 | >30 |
| L_Majors2 | FieldMatLmaj2 | Sugarloaf | 24.627298, -81.569617 | Feb 21, 2020 | Feb 21, 2020 | mature | >10 | >30 |
| L_Minors1* | FieldMatLmin1 | Sugarloaf | 24.627298, -81.569617 | Feb 21, 2020 | Feb 21, 2020 | mature | >10 | >30 |
| L_Minors2 | FieldMatLmin2 | Sugarloaf | 24.627298, -81.569617 | Feb 21, 2020 | Feb 21, 2020 | mature | >10 | >30 |
| L_Nurses1 | FieldMatLnur1 | Sugarloaf | 24.627298, -81.569617 | Feb 21, 2020 | Feb 21, 2020 | mature | >10 | >30 |
| L_Nurses2 | FieldMatLnur2 | Sugarloaf | 24.627298, -81.569617 | Feb 21, 2020 | Feb 21, 2020 | mature | >10 | >30 |
| M | FieldMatM | Conch | 24.786233, -80.896969 | Feb 22, 2020 | Feb 22, 2020 | mature | 15 | 40 |
| N_Majors1 | FieldMatNmaj1 | Duck | 24.769611, -80.914853 | Feb 23, 2020 | Feb 23, 2020 | mature | 17 | 35 |
| N_Majors2 | FieldMatNmaj2 | Duck | 24.769611, -80.914853 | Feb 23, 2020 | Feb 23, 2020 | mature | 15 | 30 |
| N_Minors1* | FieldMatNmin1 | Duck | 24.769611, -80.914853 | Feb 23, 2020 | Feb 23, 2020 | mature | 15 | 35 |
| N_Minors2 | FieldMatNmin2 | Duck | 24.769611, -80.914853 | Feb 23, 2020 | Feb 23, 2020 | mature | 30 | 30 |
| N_Nurses1 | FieldMatNnur1 | Duck | 24.769611, -80.914853 | Feb 23, 2020 | Feb 23, 2020 | mature | 30 | 26 |
| N_Nurses2 | FieldMatNnur2 | Duck | 24.769611, -80.914853 | Feb 23, 2020 | Feb 23, 2020 | mature | 20 | 18 |
| O | FieldMatO | Duck | 24.769611, -80.914853 | Feb 23, 2020 | Feb 23, 2020 | mature | 21 | 20 |
| P | FieldMatP | Duck | 24.769611, -80.914853 | Feb 23, 2020 | Feb 23, 2020 | mature | 16 | 40 |
| Q | FieldMatQ | Sugarloaf | 24.627298, -81.569617 | Feb 24, 2020 | Feb 24, 2020 | mature | 15 | >30 |
| R | FieldMatR | Sugarloaf | 24.627298, -81.569617 | Feb 24, 2020 | Feb 24, 2020 | mature | 20 | 100 |
| S | FieldMatS | Sugarloaf | 24.627298, -81.569617 | Feb 25, 2020 | Feb 25, 2020 | mature | 19.3 | 40 |
| T | FieldMatT | Sugarloaf | 24.627298, -81.569617 | Feb 25, 2020 | Feb 25, 2020 | mature | 16 | 15 |
| U | FieldMatU | Sugarloaf | 24.627298, -81.569617 | Feb 26, 2020 | Feb 26, 2020 | mature | 16.6 | 50 |
| V | FieldMatV | Conch | 24.786233, -80.896969 | Feb 27, 2020 | Feb 27, 2020 | mature | 20 | 15 |
| W | FieldMatW | Conch | 24.786233, -80.896969 | Feb 27, 2020 | Feb 27, 2020 | mature | 15 | 20 |
| X | FieldMatX | Conch | 24.786233, -80.896969 | Feb 27, 2020 | Feb 27, 2020 | mature | 21 | 21 |
| Y | FieldMatY | Conch | 24.786233, -80.896969 | Feb 27, 2020 | Feb 27, 2020 | mature | 15.3 | 22 |
| Z | FieldMatZ | Conch | 24.786233, -80.896969 | Feb 27, 2020 | Feb 27, 2020 | mature | 18 | 28 |
| cflo20.11 | FieldYoung2011 | Tea Table | 24.896367, -80.662087 | Feb 22, 2020 | Sep 28, 2020 | lab | 13.4 | 38 |
| cflo20.16 | FieldYoung2016 | Sugarloaf | 24.627298, -81.569617 | Feb 24, 2020 | Sep 28, 2020 | lab | 3.7 | 30 |
| cflo20.14 | FieldYoung2014 | Duck | 24.769611, -80.914853 | Feb 23, 2020 | Sep 28, 2020 | lab | 4.8 | 24 |
| cflo20.21 | FieldYoung2021 | Sugarloaf | 24.627298, -81.569617 | Feb 25, 2020 | Sep 29, 2020 | lab | 9.7 | 28 |
| cflo20.07 | FieldYoung2007 | Sugarloaf | 24.627298, -81.569617 | Feb 21, 2020 | Sep 29, 2020 | lab | 3.4 | 22 |
| cflo20.12 | FieldYoung2012 | Conch | 24.786233, -80.896969 | Feb 22, 2020 | Sep 29, 2020 | lab | 4.9 | 26 |
| cflo20.34 | FieldYoung2034 | Conch | 24.786233, -80.896969 | Feb 28, 2020 | Sep 29, 2020 | lab | 13.2 | 31 |
| cflo20.18 | FieldYoung2018 | Sugarloaf | 24.627298, -81.569617 | Feb 24, 2020 | Sep 29, 2020 | lab | 3.6 | 16 |
| cflo20.30 | FieldYoung2030 | Conch | 24.786233, -80.896969 | Feb 28, 2020 | Sep 29, 2020 | lab | 10.1 | 32 |
| cflo20.33 | FieldYoung2033 | Conch | 24.786233, -80.896969 | Feb 28, 2020 | Sep 29, 2020 | lab | 11.9 | 30 |
| cflo20.02 | FieldYoung2002 | Fiesta | 24.840091, -80.792447 | Feb 20, 2020 | Oct 1, 2020 | lab | 10 | 35 |
| cflo20.32 | FieldYoung2032 | Conch | 24.786233, -80.896969 | Feb 28, 2020 | Oct 1, 2020 | lab | 13.2 | 29 |
| cflo20.26 | FieldYoung2026 | Conch | 24.786233, -80.896969 | Feb 28, 2020 | Oct 1, 2020 | lab | 11.4 | 30 |
| cflo20.20 | FieldYoung2020 | Sugarloaf | 24.627298, -81.569617 | Feb 25, 2020 | Oct 1, 2020 | lab | 7.2 | 70 |
| cflo20.27 | FieldYoung2027 | Conch | 24.786233, -80.896969 | Feb 28, 2020 | Oct 1, 2020 | lab | 10 | 30 |
| cflo20.22 | FieldYoung2022 | Sugarloaf | 24.627298, -81.569617 | Feb 25, 2020 | Oct 1, 2020 | lab | 9.7 | 44 |
| cflo19.032 | LabSinInd22full | Fiesta | 24.840091, -80.792447 | Jan 20, 2019 | Sep 28, 2020 | lab2019 | 5.1 | 24 |
| cflo19.038 | LabSinInd23full | Fiesta | 24.840091, -80.792447 | Jan 20, 2019 | Sep 28, 2020 | lab2019 | 2.8 | 6 |
| cflo19.004 | LabSinInd25full | Fiesta | 24.840091, -80.792447 | Jan 17, 2019 | Sep 28, 2020 | lab2019 | 10.1 | 41 |
| cflo19.025 | LabSinInd26full | Sugarloaf | 24.627298, -81.569617 | Jan 19, 2019 | Sep 28, 2020 | lab2019 | 11 | 34 |

**SI Table 2:** Coefficient of variation by sample type. Post-hoc comparisons of gamma GLM on coefficient of variation by sample type.

| Comparison | Estimate | Std. Error | z value | Pr(>\|z\|) |  |
| --- | --- | --- | --- | --- | --- |
| ReplicaN-ReplicaL | 0.195257 | 0.036215 | 5.392 | <0.001 |  |
| Mature-ReplicaL | 0.787579 | 0.035195 | 22.377 | <0.001 |  |
| Young-ReplicaL | 0.729474 | 0.035060 | 20.807 | <0.001 |  |
| Lab2020-ReplicaL | 0.363673 | 0.036394 | 9.993 | <0.001 |  |
| Lab2019-ReplicaL | 0.197683 | 0.038955 | 5.075 | <0.001 |  |
| Nurse-ReplicaL | 0.499185 | 0.035416 | 14.095 | <0.001 |  |
| Forager-ReplicaL | 0.437798 | 0.037355 | 11.720 | <0.001 |  |
| Mature-ReplicaN | 0.592322 | 0.033084 | 17.904 | <0.001 |  |
| Young-ReplicaN | 0.534217 | 0.032939 | 16.218 | <0.001 |  |
| Lab2020-ReplicaN | 0.168416 | 0.034356 | 4.902 | <0.001 |  |
| Lab2019-ReplicaN | 0.002426 | 0.037059 | 0.065 | 1.00000 | NS |
| Nurse-ReplicaN | 0.303928 | 0.033319 | 9.122 | <0.001 |  |
| Forager-ReplicaN | 0.242541 | 0.035372 | 6.857 | <0.001 |  |
| Young-Mature | -0.058105 | 0.031815 | -1.826 | 0.60026 | NS |
| Lab2020-Mature | -0.423906 | 0.033279 | -12.738 | <0.001 |  |
| Lab2019-Mature | -0.589896 | 0.036063 | -16.357 | <0.001 |  |
| Nurse-Mature | -0.288394 | 0.032208 | -8.954 | <0.001 |  |
| Forager-Mature | -0.349781 | 0.034328 | -10.189 | <0.001 |  |
| Lab2020-Young | -0.365801 | 0.033136 | -11.039 | <0.001 |  |
| Lab2019-Young | -0.531791 | 0.035931 | -14.800 | <0.001 |  |
| Nurse-Young | -0.230289 | 0.032059 | -7.183 | <0.001 |  |
| Forager-Young | -0.291676 | 0.034189 | -8.531 | <0.001 |  |
| Lab2019-Lab2020 | -0.165989 | 0.037233 | -4.458 | <0.001 |  |
| Nurse-Lab2020 | 0.135513 | 0.033513 | 4.044 | 0.00131 |  |
| Forager-Lab2020 | 0.074125 | 0.035555 | 2.085 | 0.42312 | NS |
| Nurse-Lab2019 | 0.301502 | 0.036279 | 8.311 | <0.001 |  |
| Forager-Lab2019 | 0.240115 | 0.038173 | 6.290 | <0.001 |  |
| Forager-Nurse | -0.061387 | 0.034554 | -1.777 | 0.63467 | NS |

**SI Table 3:**
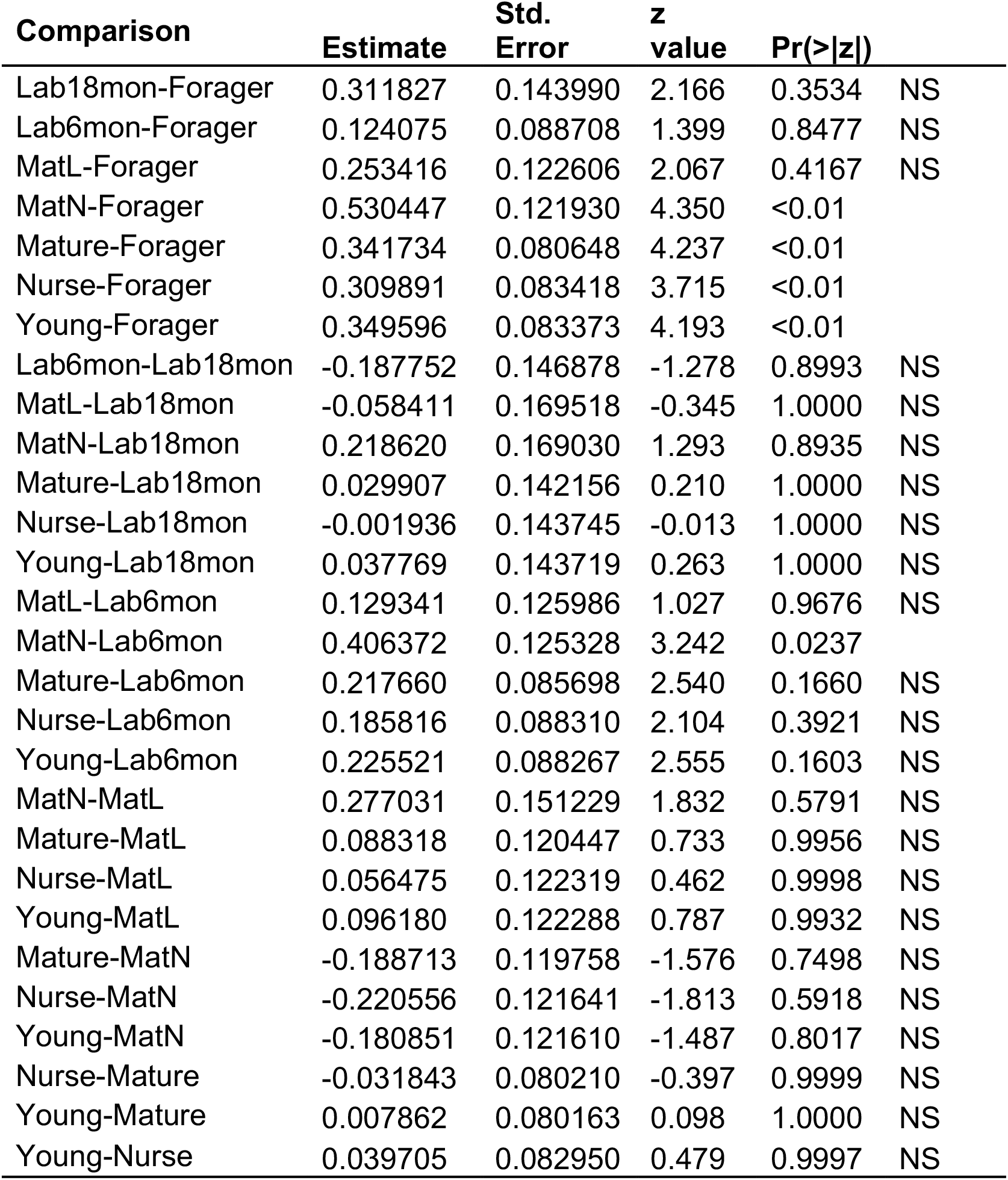
Protein number by sample type. Post hoc comparisons of negative binomial GLM on protein number explained by sample type.

**SI Table 4:** Network characteristics for all gene set enrichment analyses. The detailed results for each network are presented in SI Table 5 (for 60 most abundant proteins) and SI Table 6 (for significantly differing proteins, divided by comparison).

| Proteins used | N nodes | N edges | Avg. local clustering coefficient | Avg. node degree: | Expected number of edges: | PPI enrichment p-value: |
| --- | --- | --- | --- | --- | --- | --- |
| 60 most abundant | 41 | 32 | 0.361 | 1.56 | 8 | 2.31E-10 |
| Significant in 2/3, in any comparison | 44 | 44 | 0.312 | 2 | 13 | 2.35E-11 |
| Significant in 2/3, Nurse vs forager | 19 | 6 | 0.158 | 0.632 | 1 | 2.57E-04 |
| Significant in 2/3, Young vs mature | 10 | 4 | 0.5 | 0.8 | 1 | 1.89E-03 |
| Significant in 2/3, Field vs lab | 20 | 11 | 0.583 | 1.1 | 5 | 2.03E-02 |

**SI Table 5:**
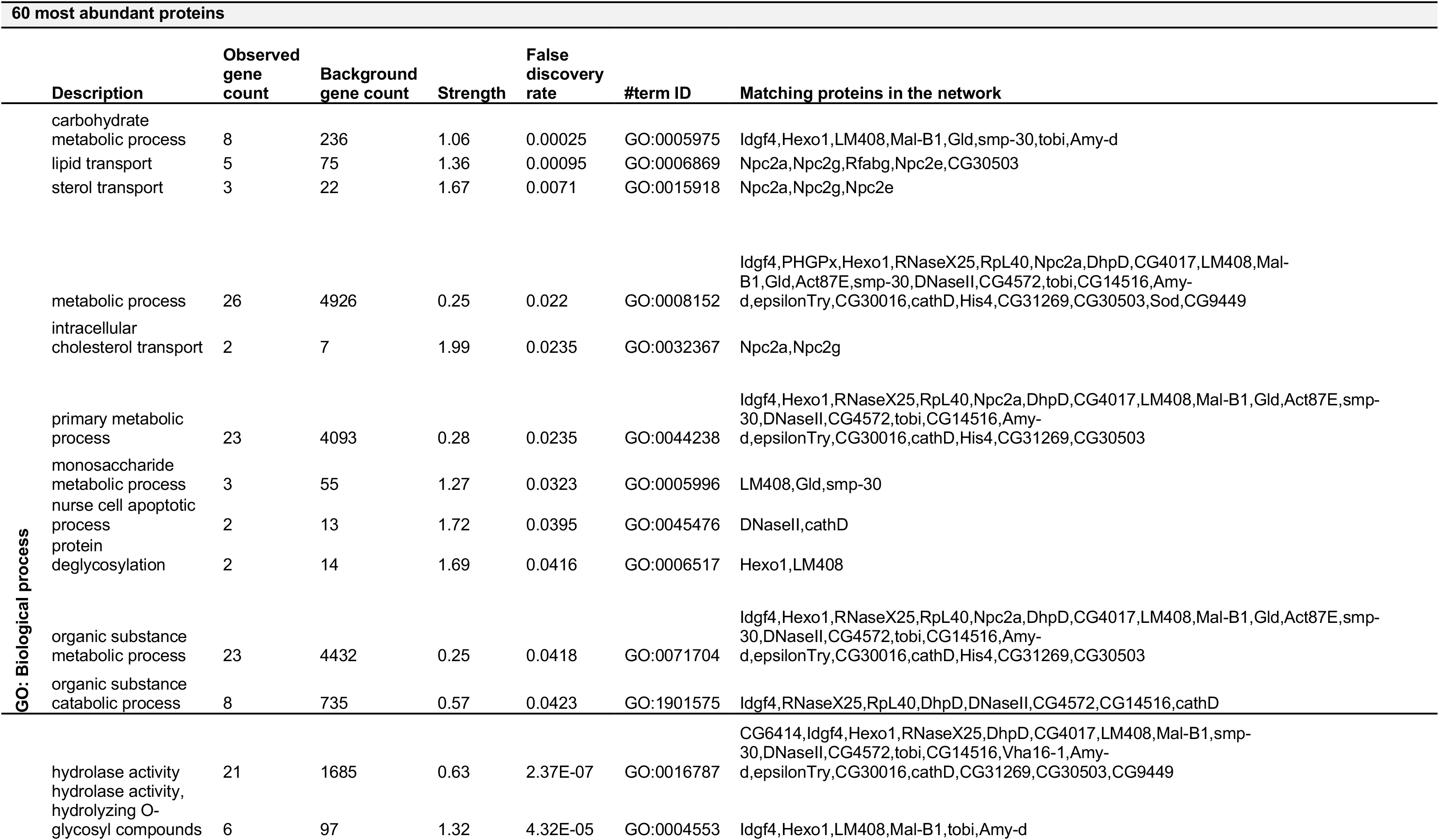
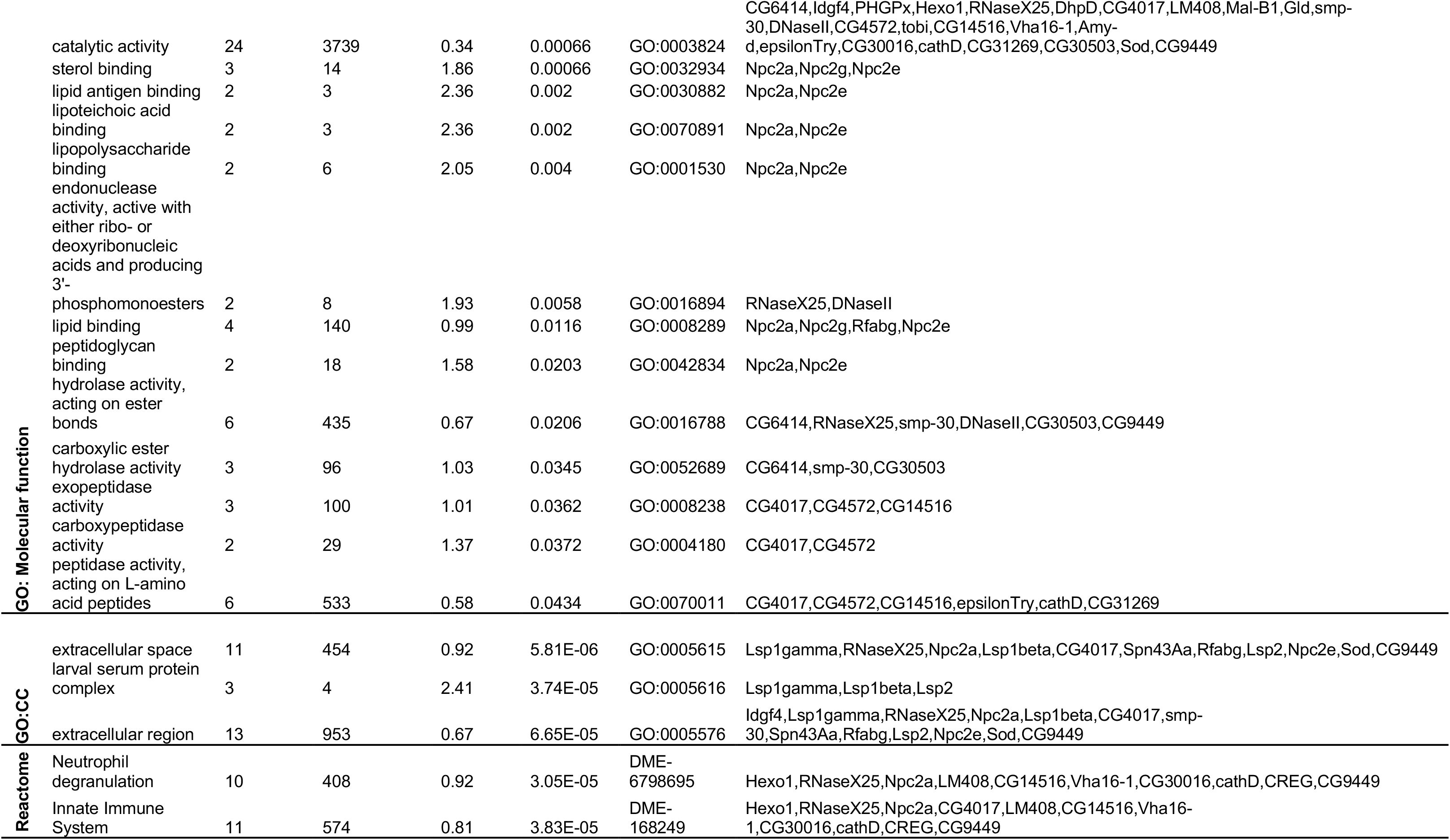
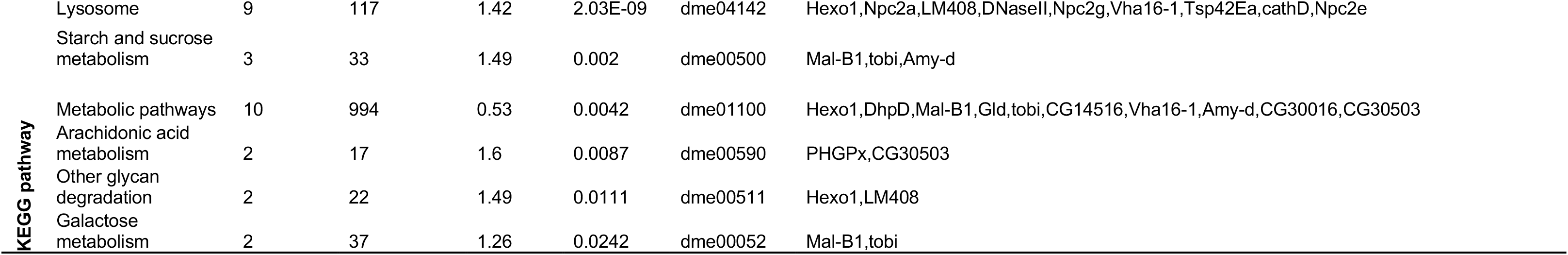
Gene set enrichment analysis for the most abundant trophallactic fluid proteins. (Next page) Gene set enrichment analysis results for the *D. melanogaster* orthologs of the 60 most abundant trophallactic fluid proteins. *Observed gene count* indicates how many proteins in the network are annotated with the term. *Background gene count* indicates how many proteins in total have this term, in this network and in the background. *Strength* describes how large the enrichment effect is: Log_10_(observed proteins / expected proteins in a random network of this size). *False Discovery Rate* describes how significant the enrichment is. P-values are corrected for multiple testing within each category using the Benjamini–Hochberg procedure.

**SI Table 6:**
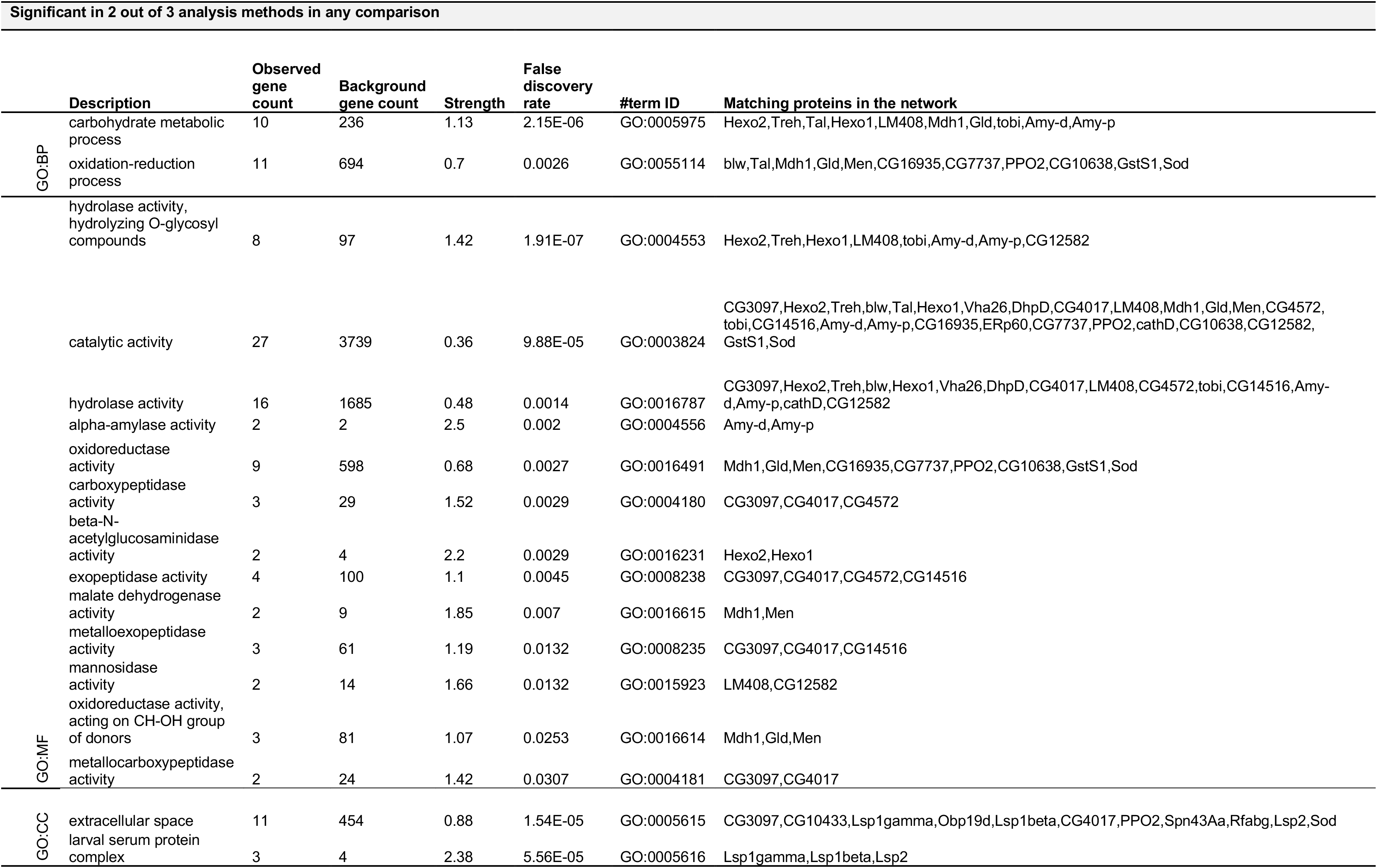
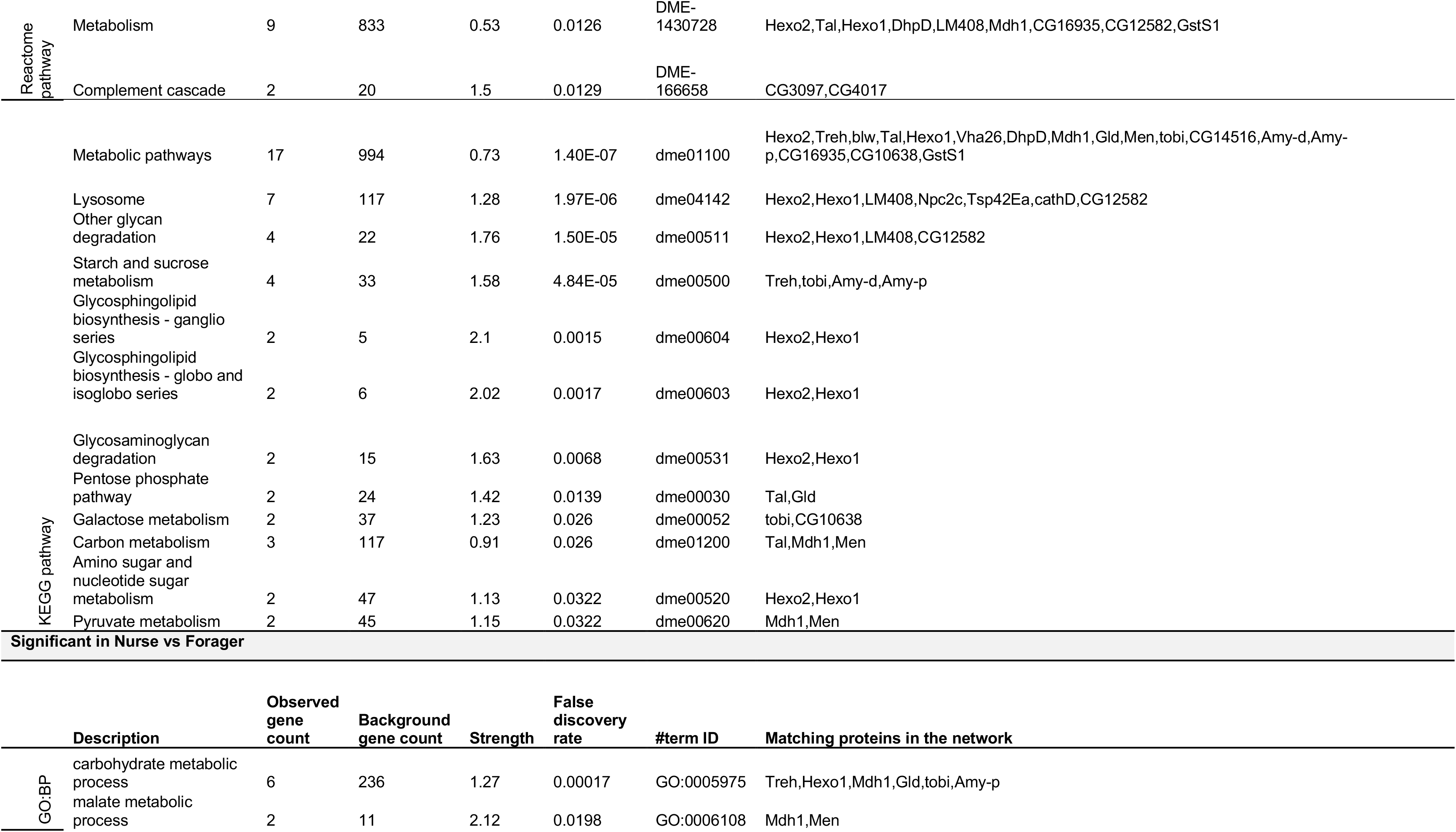
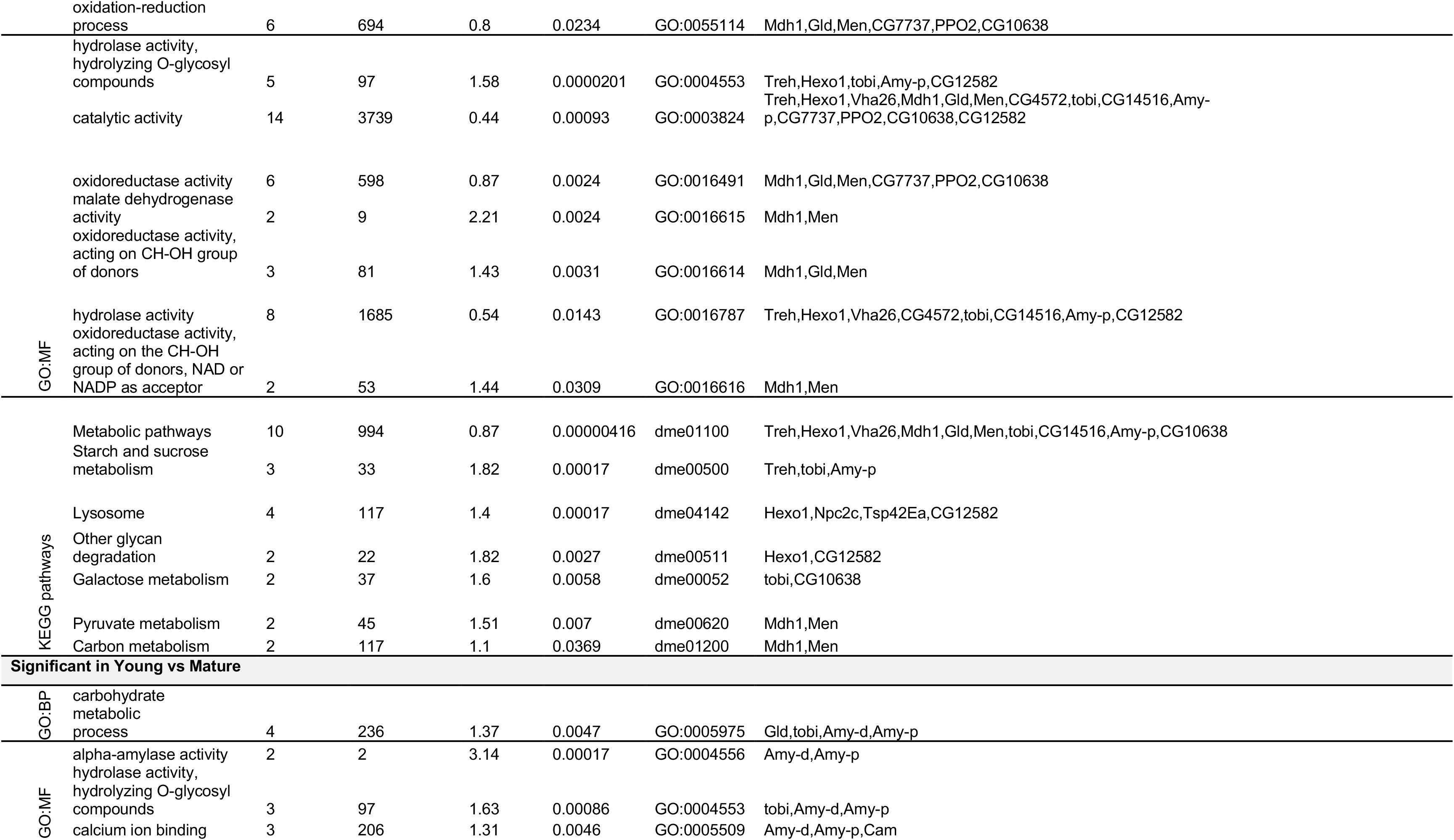
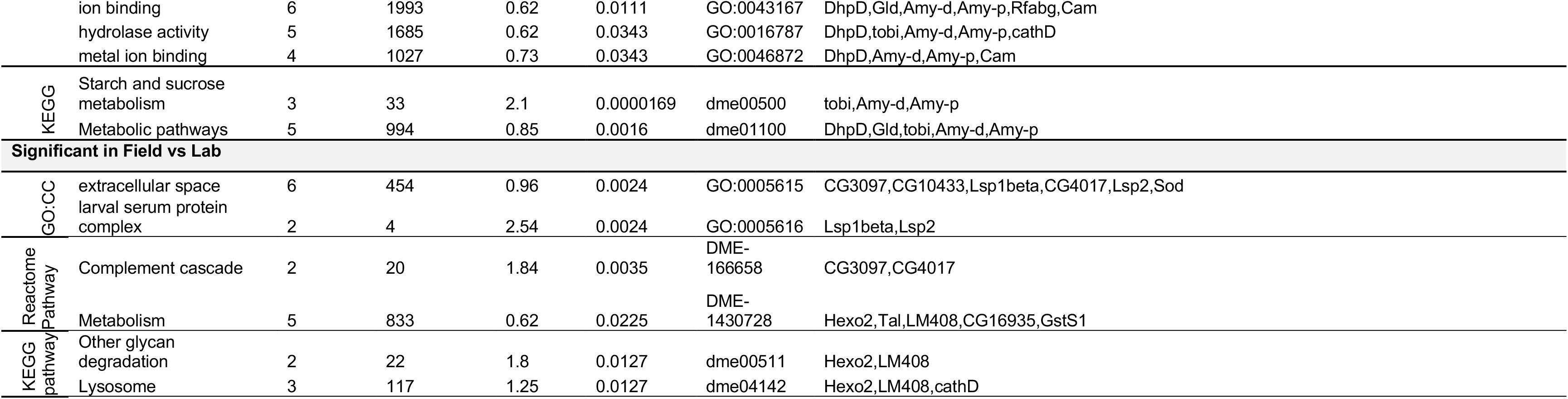
Gene set enrichment analysis for the significantly differing proteins. (Next page) Gene set enrichment analysis results for the *D. melanogaster* orthologs of the trophallactic fluid proteins significantly differing in two out of three of our statistical methods, first combined and then separately for the three main comparisons. *Observed gene count* indicates how many proteins in the network are annotated with the term. *Background gene count* indicates how many proteins in total have this term, in this network and in the background. *Strength* describes how large the enrichment effect is: log_10_(observed proteins / expected proteins in a random network of this size). *False Discovery Rate* describes how significant the enrichment is. P-values are corrected for multiple testing within each category using the Benjamini–Hochberg procedure. The significant annotations are indicated for GO: Biological process (GO:BP), GO: Molecular function (GO:MF), GO: Cellular component (GO:CC), Reactome pathways and KEGG pathways.

---

**Supplemental file 1: All 135 significantly differing proteins**

This supplemental file combines into a single sheet the results and additional information for all of the significantly differing proteins in our four comparisons (*Young vs. Mature, Nurse vs. Forager, Field vs. Lab, East vs. West*), by all of the three statistical methods (classical, empirical Bayes, machine learning). Protein accession numbers, presence in colony and individual datasets, abundance when present, fold changes by comparison and significance both by comparison and by model are shared.

**Supplemental files 2-4: Full statistical results**

These supplemental files share the full results of all the models run. For the classical frequentist models (Supplemental File 2), the imputed data is also shared. The empirical Bayes LIMMA models (Supplemental File 3), results are shared as raw output tables. Accuracy, seed, and mean feature importances for each gene are reported for each model trained for the random forest analyses in Supplemental File 4.

